# Sexual dimorphism and the multi-omic response to exercise training in rat subcutaneous white adipose tissue

**DOI:** 10.1101/2023.02.03.527012

**Authors:** Gina M. Many, James A. Sanford, Tyler J. Sagendorf, Zhenxin Hou, Pasquale Nigro, Katie Whytock, David Amar, Tiziana Caputo, Nicole R. Gay, David A. Gaul, Michael Hirshman, David Jimenez-Morales, Malene E. Lindholm, Michael J. Muehlbauer, Maria Vamvini, Bryan Bergman, Facundo M. Fernández, Laurie J. Goodyear, Eric A. Ortlund, Lauren M. Sparks, Ashley Xia, Joshua N. Adkins, Sue C. Bodine, Christopher B. Newgard, Simon Schenk, The MoTrPAC Study Group

**Author notes:** Contributed equally (Gina M. Many, James A. Sanford, Tyler J. Sagendorf).

## Abstract

Subcutaneous white adipose tissue (scWAT) is a dynamic storage and secretory organ that regulates systemic homeostasis, yet the impact of endurance exercise training and sex on its molecular landscape has not been fully established. Utilizing an integrative multi-omics approach with data generated by the Molecular Transducers of Physical Activity Consortium (MoTrPAC), we identified profound sexual dimorphism in the dynamic response of rat scWAT to endurance exercise training. Despite similar cardiorespiratory improvements, only male rats reduced whole-body adiposity, scWAT adipocyte size, and total scWAT triglyceride abundance with training. Multi-omic analyses of adipose tissue integrated with phenotypic measures identified sex-specific training responses including enrichment of mTOR signaling in females, while males displayed enhanced mitochondrial ribosome biogenesis and oxidative metabolism. Overall, this study reinforces our understanding that sex impacts scWAT biology and provides a rich resource to interrogate responses of scWAT to endurance training.

## Introduction

Subcutaneous white adipose tissue (scWAT) is a dynamic storage and secretory organ composed of lipid-storing adipocytes and numerous other cell types (e.g., immune cells, endothelial cells, mesenchymal cells, etc.)^1–3^. Through the release of a diverse array of signaling molecules such as cytokines, growth factors, lipid-derived signaling molecules, and fatty acids, scWAT impacts diverse biological processes critical for maintaining systemic health^2–12^. Indeed, numerous factors secreted by or residing in scWAT have been implicated in the development of lifestyle-related diseases such as obesity, type 2 diabetes, insulin resistance, and cardiovascular disease. Accordingly, regulation of scWAT biology—particularly in response to physiological stressors such as endurance exercise, diet, and age—is an important area for study.

Endurance exercise training (ExT) improves scWAT metabolic flexibility, lipid flux, insulin sensitivity, and immune cell polarization and expansion—factors which have been linked to the risk and severity of cardiometabolic diseases^11,13–18^. The profound effect of exercise on scWAT is illustrated by the fact that as little as 11 days of voluntary wheel running impacts the expression of thousands of genes in murine adipose tissue^11^. Despite recent advances in the field of adipocyte biology, the molecular landscape of scWAT remodeling with exercise is not fully understood. This is especially true because most ExT studies have focused on animal models of obesity. Thus, defining training adaptations in adipose tissue of lean animals may provide an important complement to such studies and provide a more balanced understanding of scWAT remodeling mechanisms.

In both humans and rodents, WAT distribution and content varies considerably between sexes^19,20^, possibly contributing to sexual dimorphism in cardiometabolic disease risk^19,21^. Although WAT is one of the most sexually dimorphic tissues^22,23^, few studies account for sex when studying exercise adaptations^11,17,18,24–26^. While many of these differences are attributable to the roles of sex hormones^19,27–29^, scWAT is sexually distinct prior to puberty, suggesting other contributors^19,27,30^. Despite these interesting and translationally relevant differences in male and female WAT, the molecular transducers orchestrating sexual dimorphism in WAT and its response to ExT remain largely unexplored.

To address these knowledge gaps, we leveraged data generated by the Molecular Transducers of Physical Activity Consortium (MoTrPAC) and a comprehensive multi-omics approach to define the temporal changes in scWAT biology in response to 1, 2, 4, or 8 weeks of treadmill ExT in male and female Fischer 344 rats^22^. This large rodent model has translational relevance for metabolic disease research due to evolutionary similarity to humans^31^, proclivity for obesity, and its development of leptin and insulin resistance in response to overnutrition^32–34^. Using this model, we identified profound sexual dimorphism in scWAT both at rest and in response to training, and highlighted candidate molecular and cellular transducers driving these differences. Provided here is a molecular compendium of the rat scWAT temporal response to progressive ExT, serving as a foundational dataset for translation to adipose-driven diseases, especially those with sex-dependent pathogenic mechanisms.

## Results

### Sexually dimorphic phenotypic responses to endurance training

Beginning at 6 months of age, male and female Fischer 344 rats underwent a progressive ExT program consisting of 5 consecutive days of treadmill running per week for 1, 2, 4, or 8 weeks at a targeted intensity of 70% VO_2_max (Fig. 1A; https://www.motrpac.org/protocols.cfm). Sedentary rats were age- and sex-matched with the 8-week-trained group. VO_2_max and NMR-derived body composition were measured at baseline and the week prior to sacrifice in sedentary, 4-week-trained, and 8-week-trained rats. Tissues were harvested from animals 48 hours after the last training session to reduce acute exercise effects. A subset of animals was randomly selected for multi-omic analyses from a larger cohort of 12–20 rats per sex per group. In animals selected for multi-omic analyses, VO_2_max (relative to lean or total body mass) increased similarly in both sexes following 8 weeks of ExT and did not change significantly in the sedentary controls (Figs. 1B, S1A). Male rats decreased body weight and whole-body fat after 8 weeks of ExT, while females maintained body fat and body weight with training (Figs. 1C, S1B–C). Sedentary females significantly increased fat and whole-body mass, whereas sedentary males only increased fat mass (Figs. 1C, S1B–C)^22^.

**Figure 1.**
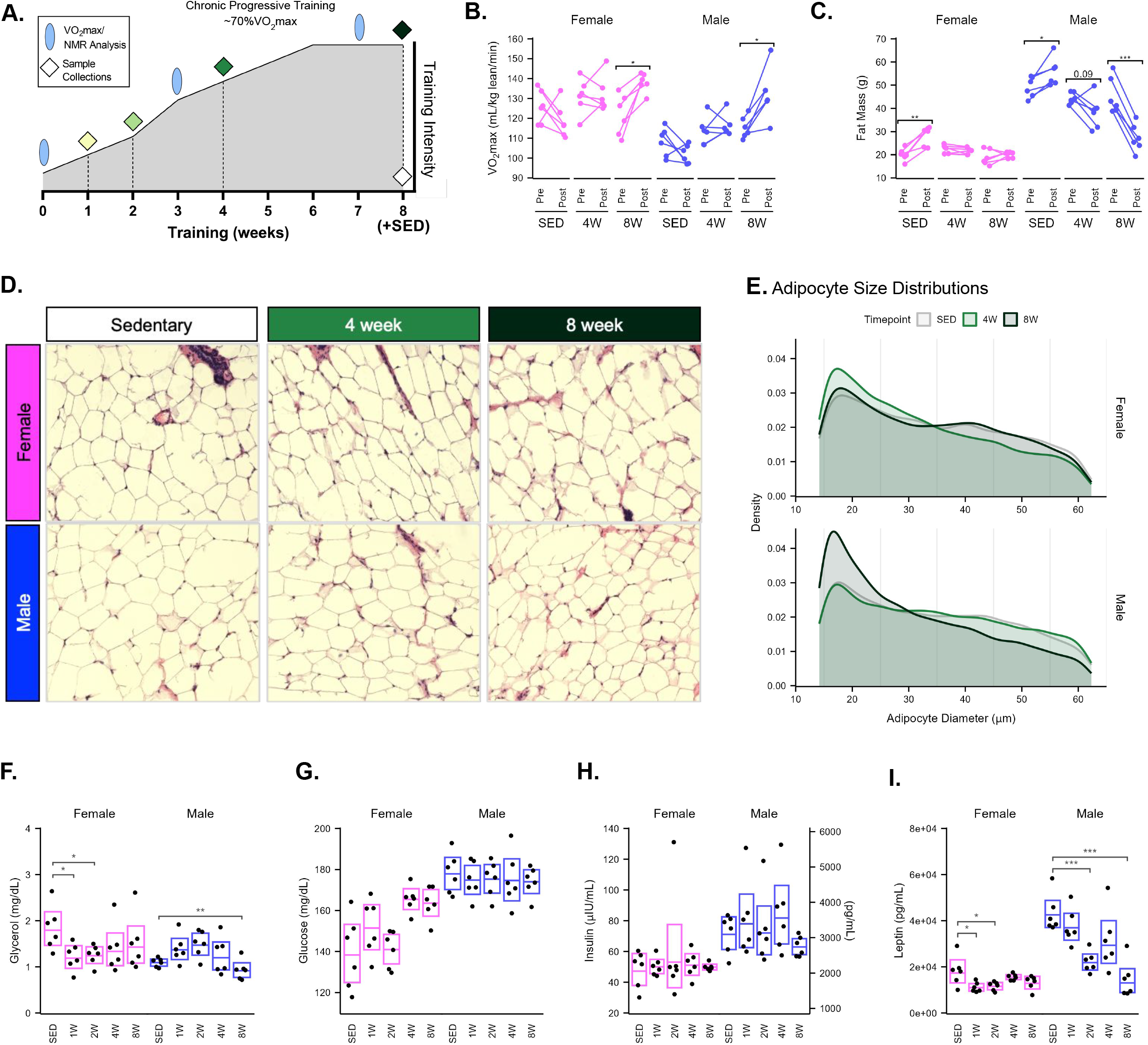
Sexually dimorphic phenotypic responses to endurance training. **A)** Diagram of exercise protocol. Fischer 344 male and female rats underwent a progressive treadmill training protocol. Tissues were collected from animals that completed 1, 2, 4, or 8 weeks of training as well as sedentary controls. Maximal oxygen consumption (VO_2_max) and nuclear magnetic resonance (NMR) body composition analysis was performed as indicated. **B)** Relative VO_2_max values (normalized to NMR lean body mass) recorded pre- and post-training from the animals selected for multi-omics analysis in the sedentary, 4-week, and 8-week-trained timepoints. **C)** Total fat mass pre and post training in sedentary, 4-week, and 8-week-trained animals, as measured by NMR. **D)** Representative histological images of scWAT from sedentary, 4-week, and 8-week-trained female and male rats. **E)** Distributions of adipocyte diameter from histological sections as measured by CellProfiler for both female and male animals. **F–I)** Measurements of circulating plasma levels of glycerol (**F**), glucose (**G**), insulin (**H**), and leptin (**I**) from rats selected for multi-omic analysis from each group. Boxes show the 95% confidence intervals of the group means, calculated from the natural-log-transformed values and then back transformed. In all plots, asterisks indicate statistical significance (*, p<0.05; **, p<0.01; ***, p<0.001); statistical tests are described in methods.

Given the changes in body composition, we examined scWAT adipocyte size following ExT (Fig. 1D). Analysis of more than 55,000 total adipocytes revealed differences in size distributions with training in both sexes (Fig. 1E). Notably, the scWAT of 4-week-trained females showed significant increases in smaller adipocytes (diameter < 20 μm) and corresponding decreases in larger adipocytes (diameter ≥ 45 μm), while 8-week-trained females returned to a distribution pattern similar to sedentary controls (Fig. 1E, S1D). In contrast, male rats had significantly higher percentages of small adipocytes in the 8-week-trained group, with significantly lower proportions of larger adipocytes (Fig. 1E, S1D), indicating a reduction in scWAT adipocyte size in males. Thus, despite similar training-induced cardiorespiratory improvements in both sexes, sexually dimorphic ExT responses occurred in overall body composition and scWAT-specific adipocyte morphology.

To further investigate sexual dimorphism in the ExT response, we measured plasma levels of common training-responsive clinical analytes. In rats selected for multi-omic analysis, plasma glycerol increased in males at 2 weeks and decreased in females at 1 and 2 weeks (p=0.006, p=0.026, and p=0.020, respectively; Fig. 1F). Non-esterified fatty acids (NEFA) decreased in plasma of male rats at week 8 (p=0.015), suggestive of triacylglycerol (TAG) mobilization from WAT in the early training response and a reduction in total fat stores with prolonged training duration in male rats (Fig. S1E). Sexual dimorphism was also apparent with regard to plasma glucose, glucagon, and insulin levels, with higher glucagon levels in females and higher glucose and insulin levels in males (Figs. 1G–H, S1F); this is consistent with previous literature showing that female rodents and humans exhibit better overall insulin sensitivity than males^19,35^. Leptin was higher in males than females at baseline and after 1 week of training; leptin levels began to decrease in males starting at 2 weeks (p<0.001) until converging with female levels at 8 weeks (Fig. 1I). While plasma leptin decreased in response to ExT in both sexes, correlation between leptin and fat loss was only observed in males (Fig. S1C, S1G). In 4-week-trained females, glucagon decreased, glucose increased, and the insulin:glucagon ratio increased (all p<0.001) (Figs. 1G, S1F, S1H). The latter is suggestive of a compensatory anabolic response to prolonged ExT in females. Plasma glucose and insulin:glucagon did not change with ExT in male rats, and HOMA-IR values did not respond to ExT in either sex (Figs. 1G, S1I). Overall, the results highlight sexual dimorphism in systemic and scWAT-specific physiological adaptations to training, several of which either promote or serve as indices of enhanced lipid mobilization in male rats. Results of these analyses are provided in Supplementary Table 1.

### Sexual dimorphism in the molecular landscape of rat scWAT

The sexually divergent clinical and physiological responses to ExT prompted a more thorough characterization of sex-specific molecular differences in scWAT. Among the 6 rats per sex per timepoint selected for multi-omic analysis, scWAT was analyzed at the transcriptomic (n=5), metabolomic (n=5), global proteomic (n=6), and phosphoproteomic (n=6) levels as described in the MoTrPAC study design^36^ using sample-level data available through the MoTrPAC Data Hub (https://motrpac-data.org/)^37^. We first compared differentially expressed molecules and pathways in sedentary rat scWAT.

Robust sex differences were observed in the abundance of molecules from all-omes—specifically, 10,336 transcripts, 4,226 proteins, 6,028 phosphosites, and 615 metabolites showed a statistically significant difference between sedentary males and females (-ome-wide false discovery rate (FDR) < 0.05), representing 20–60% of all quantified features in each dataset (Fig. 2A). Top upregulated transcripts in males included those related to lipid metabolic processes (*Car2, Acss2*) and unsaturated fatty acid synthesis (*Fads2, Fads1*) (Fig. 2A; Supplementary Table 2). Transcripts that showed higher expression in females include molecules involved in growth factor/Akt signaling (*Igf1r, Dapk1, Kit*) (Fig. 2A). At the proteomic level, top differential proteins in males were related to lipid metabolism (Acot1, Gm2a, Hadh, Acbd7), cholesterol transport (Npc2), and ketogenesis (Hmgcl, Hadh) (Fig. 2A). The scWAT of sedentary females showed higher abundances of proteins related to nuclear and cellular structural integrity (Lmnb1, Col14a1, Svil) (Fig. 2A). Notably, the oxidative stress protective factor Sod1 was upregulated at both the mRNA and protein levels in males (Fig. 2A).

**Figure 2.**
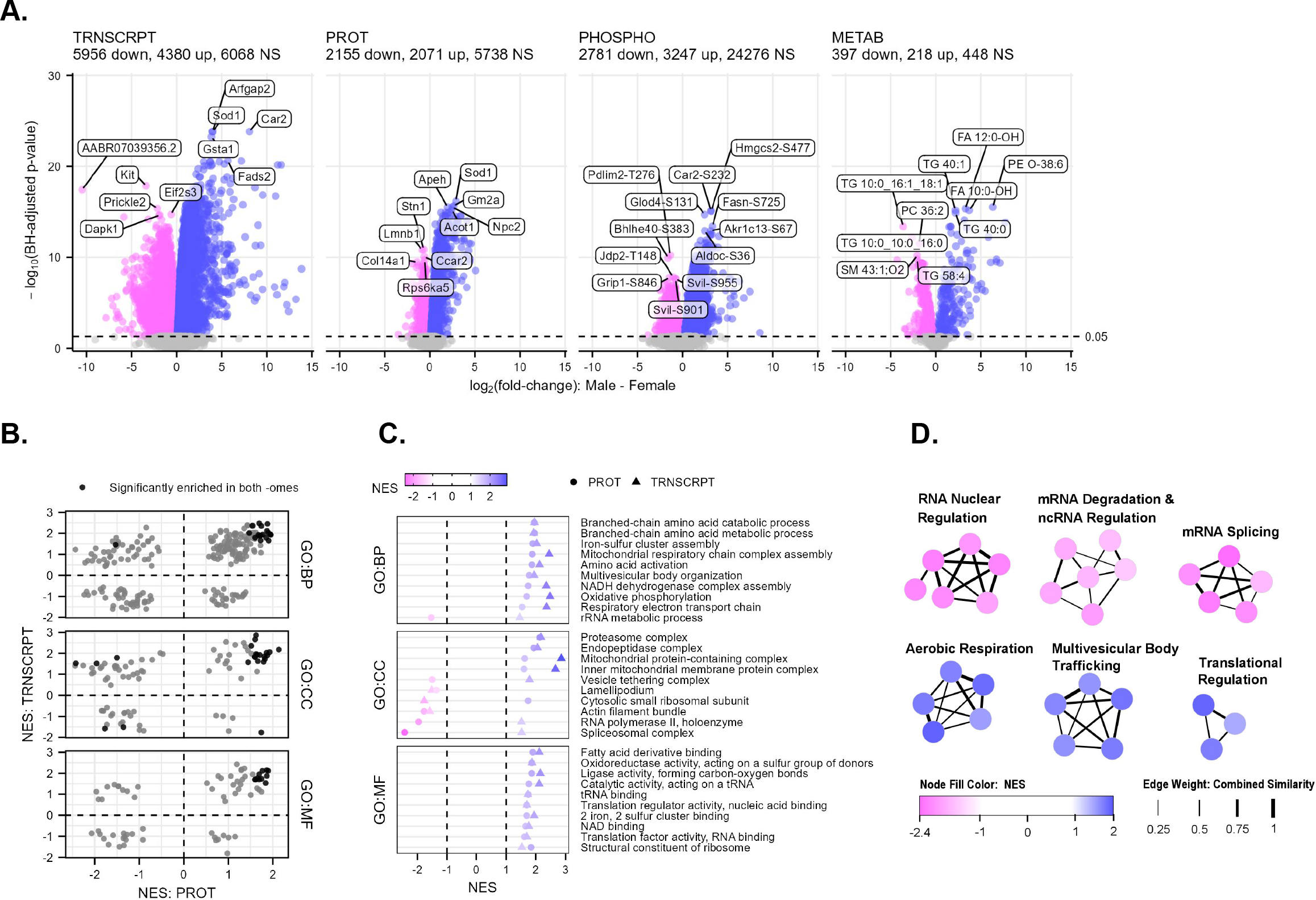
Sexual dimorphism in the molecular landscape of rat scWAT. **A)** Volcano plots displaying the magnitude and significance of changes in transcripts, proteins, phosphosites, and metabolites from the comparison of male and female sedentary controls. Top 5 or 6 features in each direction are labeled. Blue indicates that a feature was upregulated in males relative to females, while pink indicates upregulation in females relative to males. **B)** Scatterplots of the normalized enrichment scores (NES) for gene sets that were tested in both proteomics and transcriptomics separated by ontology. The black points indicate significant (BH-adjusted p-value < 0.05) enrichment in both -omes. **C)** Scatterplots of terms that were identified in (B) as being significantly enriched in both -omes in either males (blue) or females (pink). **D)** Clusters of top significantly enriched gene ontology biological processes (GO-BP) from the proteomics FGSEA comparison of sedentary male vs. female scWAT. Nodes represent individual GO-BP terms and are colored by the normalized enrichment score (NES); thickness of the edges connecting nodes relates to the proportion of genes in common between the gene sets (see methods).

Building upon these differential analyses, we performed fast gene set enrichment analysis (FGSEA)^38^ to test for enrichment of Gene Ontology (GO) terms (biological processes (BP), molecular functions (MF), and cellular components (CC)) (Supplementary Table 3) and evaluated pathway-level consistency between transcriptomic and proteomic comparisons of sedentary male and female scWAT (Fig. 2B). GO terms related to oxidative metabolism, mitochondrial complex assembly, branched-chain amino acid (BCAA) catabolism, proteasome activation, and ribosome subunits were consistently enriched in males at both the transcriptomic and proteomic levels (Fig. 2C). In females, terms including lamellipodium and actin filament bundles were enriched in both -omes (Fig. 2C). When probing the top proteomic networks displaying sexually-distinct enrichment, sedentary males showed enrichment of terms related to aerobic metabolism, multivesicular trafficking (including vesicle ubiquitination-dependent catabolism and sorting), and translational regulation (Fig. 2D). Females displayed upregulation of networks related to mRNA nuclear regulation (including export), mRNA splicing, and mRNA degradation and ncRNA regulatory processes (5.8S RNA maturation and snoRNA metabolism) (Fig. 2D).

FGSEA of RefMet metabolite subclasses^39^ also uncovered sexual dimorphism, with enrichment of amino acid and acyl-CoA species in sedentary males. Female scWAT displayed enrichment of TAG, primarily driven by several long-chain (>40 carbons) TAG species (Supplementary Table 3). Such metabolomic differences are consistent with enrichment of mitochondrial and amino acid pathways at the transcriptomic and proteomic levels in males. Taken together, the multi-omic measures demonstrate remarkable differences in scWAT between sedentary male and female F344 rats at the transcriptomic, proteomic, and metabolomic levels, and provide a new foundation for understanding sexual dimorphism in the scWAT ExT response.

### Sex-specific multi-omic scWAT adaptations to ExT

We next analyzed the sample-level datasets to identify differentially expressed features at each training timepoint for each sex (e.g., 1-week females versus SED females). As was the case with several physiologic phenotype responses, overt sexual dimorphism was observed in the multi-omic response to ExT (Fig. 3A–B; Supplementary Table 4). In general, female scWAT displayed a comparable total number of differential features at all 4 ExT timepoints, whereas the magnitude of the scWAT multi-omic response increased throughout training in males, resulting in a greater number of differentially regulated features at the later ExT timepoints in males (Fig. 3A–B).

**Figure 3.**
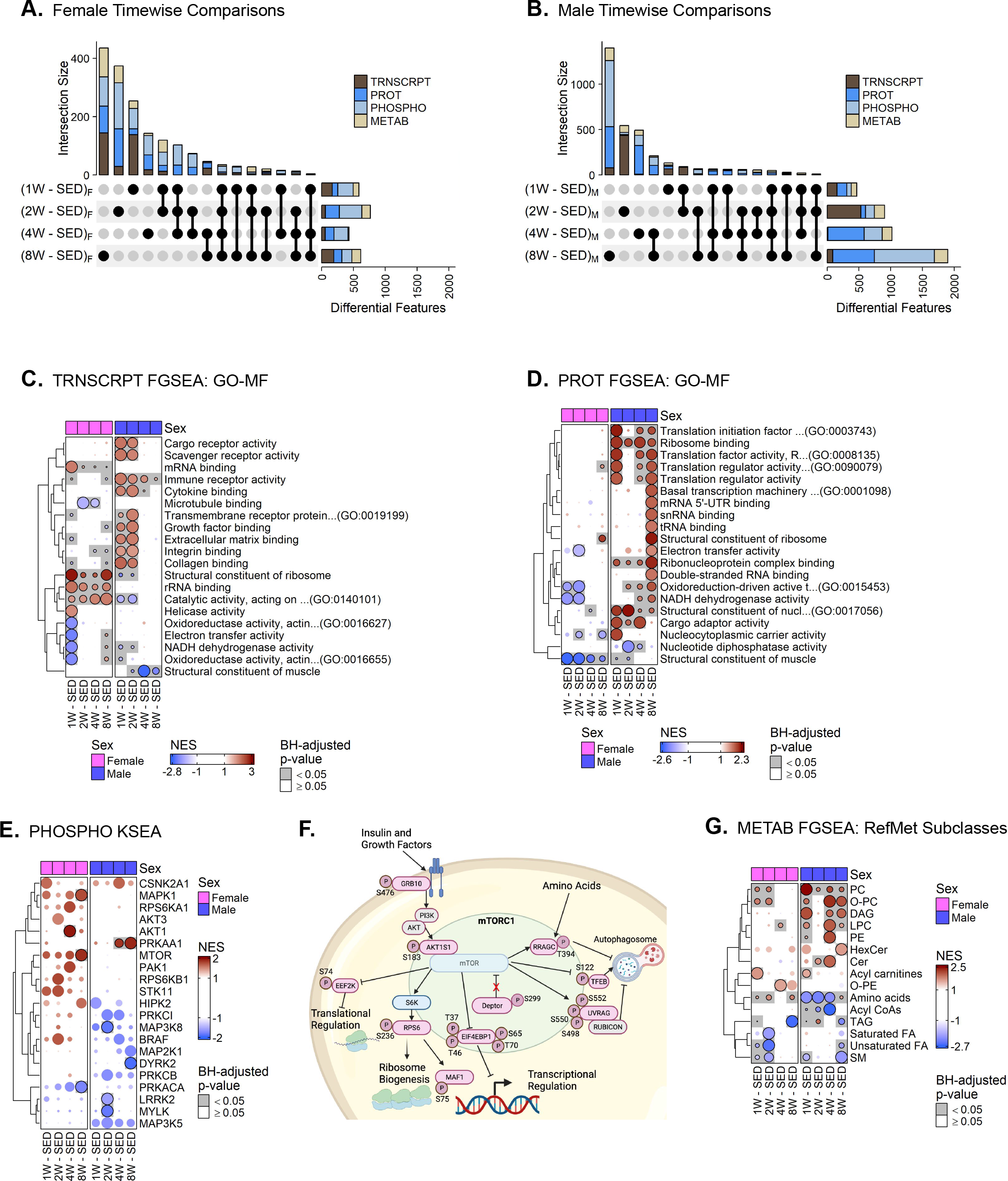
Sex-specific multi-omic scWAT adaptations to ExT. **A–B)** UpSet plots of statistically significant (FDR<0.05) transcripts, proteins, phosphosites, and metabolites from each comparison between trained and sex-matched sedentary control female (**A**) and male (**B**) rats. **C–D)** Top molecular functions (MF) from the Gene Ontology database that are most significantly enriched in any of the 8 comparisons from the transcriptomics (**C**) or proteomics (**D**) FGSEA results. Circles are colored by the NES and scaled by row so that the most significant comparison is of maximum area. **E)** Inferred activity of the indicated kinases in each of the trained groups versus sex-matched sedentary controls from Kinase–Substrate Enrichment Analysis (KSEA) of the phosphoproteomics differential analysis results. **F)** Schematic of significant phosphorylation changes driving MTOR enrichment in 8-week-trained females; phosphosites driving enrichment are colored pink. **G)** RefMet chemical subclasses that are significantly enriched in at least one of the 8 comparisons according to the metabolomics FGSEA results.

#### Transcriptomics

ExT induced distinct temporal patterns of differentially regulated transcripts—representing a combined 788 and 474 differentially regulated transcripts at discrete timepoints in males and females, respectively (FDR < 0.05). The most robust responses occurred in 1- and 8-week-trained females (167 and 193 transcripts, respectively) and 2-week-trained males (529 transcripts) (Figs. 3A–B, S2A). Six transcripts displayed differential regulation at all training timepoints in females. Notably, *Grb14*, a negative modulator of insulin signaling, was downregulated with ExT while *Olah*, a thioesterase involved in medium chain fatty acid synthesis, and *Hmgcs2*, the rate-limiting enzyme of cholesterol synthesis, were increased at all training timepoints in females (Supplementary Table 4). The only transcript that displayed differential expression at all training timepoints in males was the hypoxia-inducible carbonic anhydrase (*Ca12*)^40^, which was downregulated. FGSEA further highlighted sexual dimorphism in the progressive transcriptomic response to ExT (Figs. 3C, S3A–B). After 1 and 2 weeks of ExT, male scWAT was positively enriched for terms related to immune receptor activity and binding of cytokines, growth factors, extracellular matrix, integrins and collagens (Fig. 3C). These pathway enrichments were driven by genes involved in tissue stress (*Hif1a, Tlr4, C4b, Nlrp3*), tissue remodeling (*Itgb1, Timp3, Lifr, Pten, Fgfr1*), and angiogenesis (*Kdr, Flt1, Pdgfb, Vegfa*) (Figs. S3A, 3C; Supplementary Table 5). Enrichment of the GO-MF cytokine binding and immune receptor activity pathways was driven by transcripts related to Th17 cell activation (*Cd4, Il17ra, Tgfbr2, Il6st, Ctsl*) in male scWAT at 2 weeks. Females displayed early enrichment of terms related to transcription/translational regulation that generally peaked at 1 week and remained enriched throughout training (Fig. 3C; S3A–B). Both males and females displayed an early (1 week) adaptive immune response that was attenuated at later timepoints in females, but remained positively enriched in males throughout ExT (Fig. S3A). Terms related to oxidative phosphorylation (OXPHOS) activity decreased in both sexes at 1 week but recovered by 2 weeks in males and 4 weeks in females (Fig. 3C). In 8-week-trained rats, terms related to the adaptive immune system, immune receptor activity, azurophil granules, and vacuolar lumen were enriched in males, whereas transcription and translation regulatory processes, aerobic respiration, and azurophil granules were enriched in 8-week-trained females (Figs. 3C, S3B). Complete results of these analyses are provided in Supplementary Tables 4 and 5.

#### Proteomics

There was a robust change in the scWAT proteome in male rats that increased with training duration (654 differentially expressed proteins at 8 weeks), while the response in females was much smaller, with most differences occurring after 2 weeks (221 differential proteins) (Figs. 3A–B, S2B). In males, 18 proteins were differentially regulated at all timepoints, most of which were related to mitochondrial function (Maip, Atp6v0d1, Immt), solute transport (Slc25a11, Slc25a3, Slc25a15), and vesicle transport (Tmed2, Sec61a1). Lrpprc, a protein that regulates transcription of mitochondrial genes to promote lipid oxidation^41^, was increased throughout training in males (Supplementary Table 4). Slc25a15, a mitochondrial ornithine carrier that promotes arginine synthesis, was also elevated throughout training in male scWAT. Arginine promotes lipolysis partially through Ampk activation^42^. FGSEA revealed a robust increase in GO terms related to OXPHOS and mitochondria-specific ribosome biogenesis in male scWAT starting at 2 weeks of ExT (Figs. 3D, S3C; Supplementary Table 5). Males also displayed enrichment of terms related to vesicle transport and ribosome activity and biogenesis throughout training (Figs. 3D, S3C–D). Females displayed differential regulation of 5 proteins at all 4 timepoints. Consistent with transcriptomic profiling, the insulin signaling repressor Grb14^43^ was downregulated in females at all 4 timepoints. In contrast to males, female scWAT displayed negative enrichment of terms related to mitochondrial processes at 1 and 2 weeks, with modest enrichment of mitochondrial-related terms at 8 weeks (Fig. S3C–D). Both sexes displayed enrichment of terms related to transcription and translation at 8 weeks, with a more robust proteomic than transcriptomic response in males (Figs. 3D, S3C–D).

#### Phosphoproteomics

Several hundred protein phosphorylation sites were differentially regulated with ExT, with sexually dimorphic temporal patterns similar to proteomic ExT responses. In general, males displayed a more robust phosphoproteomic response following 4 and 8 weeks of ExT (Figs. 3A–B, S2C). Females modulated 14 phosphoproteins at all 4 training timepoints, including decreased phosphorylation of proteins related to inflammatory responses (Pde4 a–b, Ankrd2, Lrrfip1) and Camk2b, a negative regulator of adipocyte insulin signaling^44^. Males exhibited consistent modulation of the phosphorylation of 15 proteins throughout ExT, including increased phosphorylation of proteins regulating vesicle transport (Htt-S621, Klc1-S521/S524, Uhrf1bp1l-S418) and autophagy (Bnip3-T66, Ulf1-S458). To allow for inference of kinase activity based on changes in phosphorylation of known substrates, we utilized curated kinase–substrate relationship data from PhosphoSitePlus (PSP)^45^ to conduct kinase–substrate enrichment analysis (KSEA) (Fig. 3E). The results of this analysis showed decreased activity of MAP3K8, LRRK2, and MYLK kinases in 2-week-trained males (Fig. 3E) whereas PRKAA1 (AMPK1ɑ1) activity increased in 4- and 8-week-trained males. DYRK2 activity decreased at 8 weeks in males (Fig. 3E), which was largely driven by changes in growth factor-responsive CARHSP1 phosphorylation (S30, S32, S41)^46^. In females, AKT1 activity increased at 4 weeks before subsiding by week 8 when MAPK1 and mTOR kinase activity increased. Increases in mTOR kinase activity were driven by phosphorylation of RAGC at the insulin-dependent T394 phosphosite, EIF4EBP1 at S65 and T70, AKT1S1 (PRAS40) at S183, and MAF1 at S68 and S75 (Supplementary Table 5); a graphical illustration of phosphosites driving MTOR enrichment is presented in Figure 3F. Phosphorylation of EIF4EBP1 permits MTOR-mediated transcription^47^, while phosphorylation of AKT1S1 and MAF1 permits insulin-mediated MTOR activation via AKT^48^, thus suggestive of a training-induced enrichment of mTORC1 activity in female scWAT. In females, phosphosites driving AKT1 enrichment included the AKT-dependent inactivation site of catabolic FOXO3 (S253) and the fatty acid synthesis protein ATP citrate lyase (ACLY) activation sites, S425 and T447^49^. Overall enrichment in AKT and MTOR signaling in female scWAT suggests a female-specific mechanism to promote scWAT lipid storage and preserve scWAT fat stores in response to ExT^50^.

#### Metabolomics

We employed a comprehensive suite of metabolomics technologies to characterize changes in the scWAT metabolome/lipidome with ExT, including non-targeted methods as well as quantitative targeted methods focused on clusters of chemically related analytes such as amino acids, acylcarnitines, and nucleotides^36^. Overall, the scWAT metabolome displayed the least sexual dimorphism in the omic response to ExT, with the most differentially regulated features appearing at 2 and 8 weeks in both sexes (2 weeks: 135 and 161 differential features in females and males, respectively; 8 weeks: 139 and 211) (Figs. 3A–B, S2D). The 4-week-trained timepoint displayed the most sexual dimorphism with 154 differentially regulated metabolites in males and only 20 in females. In both sexes, FGSEA of RefMet metabolite subclasses^39^ revealed positive enrichment of phosphatidylcholine (PC), ether phosphatidylcholines (O-PC), and acylcarnitine species in 1-week-trained rats, followed by negative enrichment of TAGs at 8 weeks (Fig. 3G). Females displayed a prolonged early enrichment of PC and O-PC phospholipid species, along with a negative enrichment of sphingomyelin (SM) and unsaturated fatty acids (FA) at 1 and 2 weeks. Males showed positive enrichment of PC, diacylglycerols (DAG), and lysophosphatidylcholines (LPC) at 1, 4, and 8 weeks of ExT (Fig. 3G). Males additionally displayed an increase in phosphatidylethanolamine (PE) at 4 weeks that subsided by week 8 (Fig. 3G). PE and PC are the major phospholipid species present in the phospholipid monolayer of the lipid droplet (LD). The increase of PC relative to PE in 8-week-trained males is consistent with our observation of reduced adipocyte size, which suggests that males halt LD formation after 8 weeks, supported by phenotypic changes in adipocyte size distribution and total fat mass^51,52^. Notably, acyl-CoAs, which were elevated in the scWAT of sedentary males, decreased in 4-week-trained males (Figs. 3G, S4A), further supporting increased lipid utilization with ExT in males. Females displayed positive enrichment of amino acids at 1, 2, and 8 weeks, whereas amino acid species decreased in males through the first 4 weeks of training before they rose sharply at 8 weeks (Figs. 3G, S4B); this is perhaps suggestive of mobilization of amino acids from protein as lipid stores are depleted in males in response to prolonged training. This interpretation is also consistent with the surge in mono- and di-phosphonucleotides, reduction in tri-phosphonucleotides, and increased AMP:ATP ratio corresponding to a rebound in scWAT fatty acyl-CoA abundance at 8 weeks in males (Fig. S4C).

#### Sexual dimorphism in the temporal response to ExT

Informed by the strong sex differences in scWAT of sedentary animals, we sought to more directly examine the sexual dimorphism in response to progressive ExT. To this end, we compared the male ExT response to the female response at each trained timepoint (e.g., (Males.1W - Males.SED) - (Females.1W - Females.SED), etc.), which highlights differences in the magnitude and/or direction of training responses between sexes. These analyses revealed that at the transcriptomic level, pathways related to immune receptor activation, GTPase activation, and transcriptional/translational regulation showed the most divergent responses between the sexes. (Fig. S5A–B). At the proteomic level, there were strong sex differences in exercise-induced changes to pathways related to mitochondrial activity, vesicle transport, and transcriptional/translational processes (Fig. S5C–D). The strongest phosphoproteomic differences with prolonged ExT were increased MTOR kinase activity in 8-week-trained females and increased PRKAA1 (AMPKɑ1) activity in 4- and 8-week-trained males (Fig. S5E). The metabolomic response to ExT displayed sexual divergence in amino acid, acyl-CoA, DAG, PC, O-PC, and LPC species abundance (Fig. S5F).

### Integrative phenotypic-omic responses to ExT

To integrate -omics measurements, we used weighted gene co-expression network analysis (WGCNA) to generate distinct modules for metabolomics/lipidomics, proteomics, and transcriptomics datasets (Fig. S6A–C). We then assessed correlations of the module eigenfeatures (MEs)—the principal eigenvector of each module—with select clinical plasma analytes and other phenotypic markers (Fig. 4A), as well as correlations between metabolomics and other -omics MEs (Fig. 4B). Afterwards, over-representation analysis (ORA)^53^ was performed to test for localization of Gene Ontology terms or RefMet chemical subclasses in each module.

**Figure 4.**
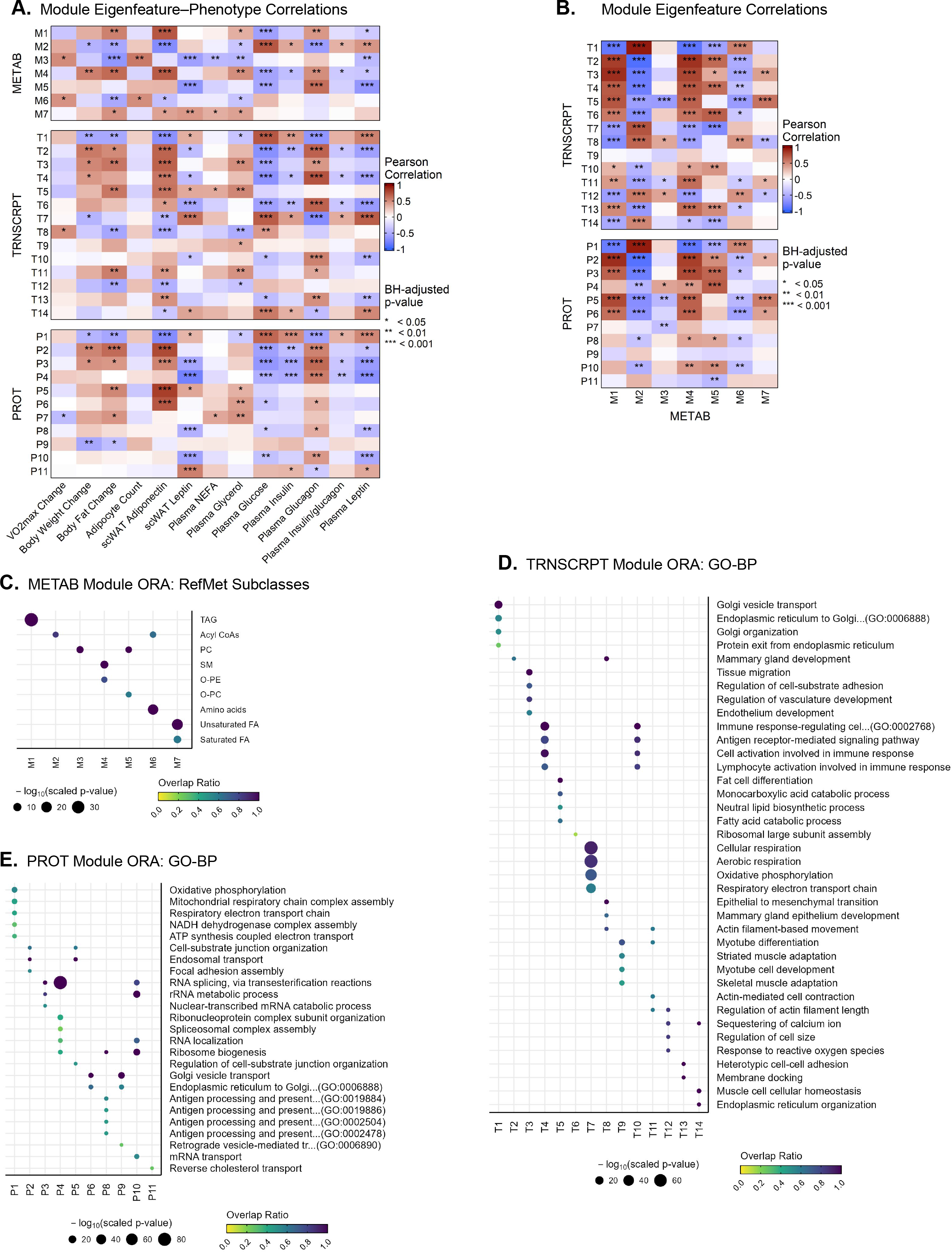
Integrative phenotypic–omic responses to ExT. **A)** Heatmaps of correlations between module eigenfeatures (MEs) and clinical measurements. **B)** Heatmaps of correlations between the metabolomics MEs and the MEs of the other -omes. **C)** All over-represented RefMet chemical subclasses in each metabolomics WGCNA module. **D–E)** Top over-represented GO-BP terms in each transcriptomics (**D**) or proteomics (**E**) WGCNA module.

#### Metabolomics

The ME of the largest metabolomics/lipidomics module, M1, was higher in females throughout training (Fig. S6A); M1 was over-represented by long chain (>48) TAG species (Fig. 4C). Like the other two modules with higher MEs in females (M4, over-represented by SM and O-PE species; M5, over-represented by PC and O-PC species), the M1 ME correlated positively with scWAT adiponectin and fat mass, and negatively correlated with plasma glucose levels (Figs. 4A, 4C, S6A). The second largest module, M2, was over-represented by acyl-CoA species (Fig. 4C); its ME was higher in males and negatively correlated with scWAT adiponectin and changes in body fat (fat loss) (Fig. 4A). M3 contained predominantly 38–40 carbon PC species (Fig. 4C), and its ME increased at 8 weeks in males (Fig. S6A). The MEs of M3 and the amino-acid-containing M6 module correlated positively with scWAT adipocyte count per field (reduced adipocyte size) and negatively with body fat changes (Fig. 4A, 4C), suggesting correlation with adipocyte lipolysis.

#### Transcriptomics

T1 was the largest transcriptomics module and its ME was highest in males, slightly increasing with training in females (Fig. S6B). T1 was over-represented by terms relating to vesicle export and recycling processes, including autophagosome and protease terms (Fig. 4D; Supplementary Table 6). M1 also contained terms relating to mitochondrial complex assembly and fatty acid catabolic processes (Supplementary Table 6); its ME negatively correlated with the ME of the TAG-localizing M1 metabolomics module (Fig. 4B), indicative of an inverse relationship between the aforementioned processes and scWAT TAG storage. The T7 ME was also higher in males and the module was over-represented by aerobic respiration terms (Figs. S6B, 4D). This ME was negatively correlated with the TAG-localizing M1 module, and positively correlated with the ME of the acyl-CoA-containing M2 ME (Figs. S6B, 4B). The MEs of T2–T4 were higher in females and correlated with scWAT adiponectin protein abundance (Figs. S6B, 4A). T3 was over-represented by terms related to developmental processes, including those containing classic adipogenic progenitor transcripts (*Cd34*, *Dpp4*, *Icam1*, *Pdgfrb*)^1^, alluding to elevated adipogenic potential in female scWAT (Fig. 4D). This ME correlated with plasma glucagon and the TAG-localizing M1 ME (Fig. 4A–B), suggestive of a possible lipid-sparing role for these transcripts. The T5 ME also correlated with the ME of the TAG-localizing M1 and remained relatively stable in females, increasing and then decreasing at 8 weeks in males, which resulted in elevated ME abundance in females relative to males at 8 weeks (Figs. 4B, S6B). ORA of T5 revealed localization of transcripts involved in lipid storage and fat cell differentiation (*Fabp4*, *Lipe*, *Cebpa*, *Plin1*, *Pparg*, *Adipoq*, *Lpl, Slc2a4 [Glut4]*) (Supplementary Table 6), suggestive of an adipogenic module. T4 and T10 were over-represented by immune signaling terms, most notably lymphocyte activation, that increased only in males with training (T4 transcripts: *Btk, Cd28*, *Cd8a*, *Ctla4, Klrd1, Klrk1, Rela [Nfkb], Themis*; T10 transcripts: *Ptprc* [*Cd45*], *Cd19*, *Cd22*, *Foxp3*, *Il21, RT1-DOa*) (Figs. 4D, S6B). Interestingly, both MEs negatively correlated with scWAT and plasma leptin levels (Fig. 4A), potentially suggestive of lymphocyte-mediated lipolysis.

#### Proteomics

The ME of the largest proteomics module, P1, was much higher in males and increased in both sexes with training (Fig. S6C). P1 was dominated by terms relating to mitochondrial aerobic respiration/OXPHOS (Fig. 4E). The P1 ME was negatively correlated with fat and body mass, the TAG-containing M1 ME and scWAT adiponectin levels (Fig. 4A–B), illustrating a negative association between lipid oxidation and scWAT TAG abundance. Interestingly, P1 also contained proteins related to peroxisomal tethering and biogenesis (Pex6, Pex7, Pex10, Pex13, Pex19), proteasomal proteins, and activation of the innate immune system (e.g., azurophil granule) (Supplementary Table 6). Further supportive of a metabolic-regulatory role of P1 proteins was a strong correlation between the P1 ME and the amino acid-dominant M6 ME (Fig. 4B). The MEs of the next largest modules, P2–P4, were more abundant in females (Fig. S6C). These modules were over-represented by terms related to endosomal transport (P2), acetyltransferase activity and histone acetyltransferase complexes (P3), and RNA regulatory processes (P3–P4) (Fig. 4E; Supplementary Table 6); their MEs correlated with body fat and scWAT adiponectin levels (P2–P3) and negatively correlated with plasma glucose, insulin, and leptin (P2–P4) (Fig. 4A). These modules were positively associated with M4 and M5 MEs.

### Integrative -omics reveals sexual dimorphism in scWAT mitochondrial metabolism and lipid recycling with ExT

FGSEA of the MitoCarta3.0 database^54^ was leveraged to provide deeper insight into sexual dimorphism of scWAT energy metabolism. Consistent with our previous FGSEA results, sedentary males showed strong enrichment of MitoCarta terms compared to females (Figs. 2C, 5A), including increased abundance of proteins involved in fatty acid oxidation, BCAA metabolism, and OXPHOS. Notably, no MitoCarta pathways were significantly enriched in sedentary females over males, further emphasizing baseline sexual dimorphism in scWAT metabolism. Moreover, males showed enhanced enrichment of catabolic pathways such as BCAA metabolism, carbohydrate metabolism, fatty acid oxidation, proteases, and gluconeogenesis over the training time course that were most apparent at 8 weeks; meanwhile, females exhibited enrichment in BCAA catabolism at 4 weeks that attenuated at week 8 with no or very modest increases in other catabolic pathways (Fig. 5B). Other MitoCarta terms relating to protein metabolism were co-induced with prolonged duration of ExT in female and male scWAT, including protein import and sorting, protein homeostasis, and chaperones (Fig. 5B). Despite these changes in multiple pathways relating to mitochondrial metabolism, we found that markers of mitochondrial abundance as assessed by mtDNA content, percentage of RNA-Seq reads mapping to mitochondrial genes, and cardiolipin CL(72:8) abundance were unaffected by training in either sex (Figs. 5C, S7A). These results suggest that scWAT from males is qualitatively poised for oxidative metabolism in the sedentary state and further adapts with ExT to favor mitochondrial metabolism and fuel catabolism due to changes in mitochondrial activity rather than biogenesis.

**Figure 5.**
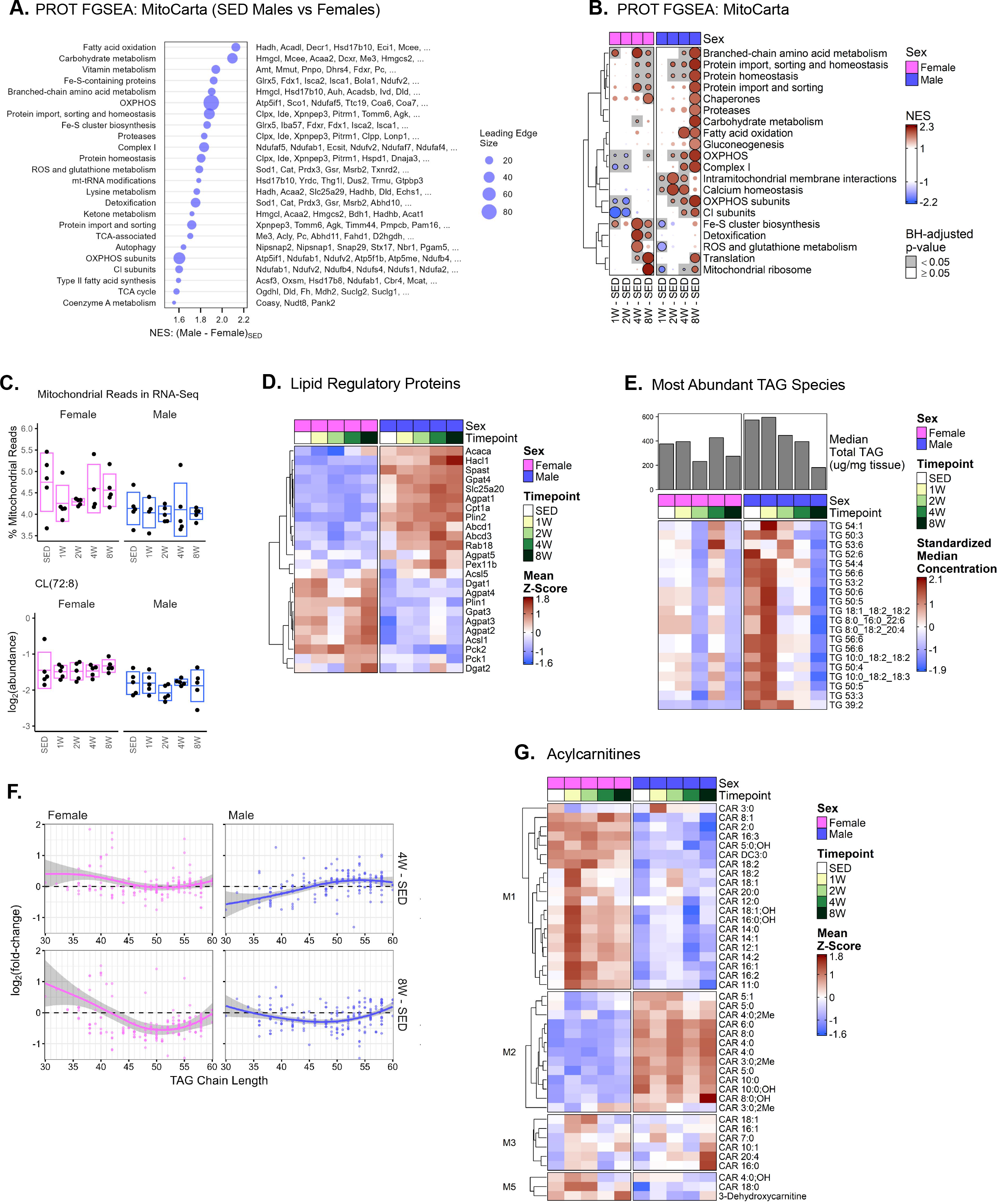
Integrative -omics reveals sexual dimorphism in scWAT mitochondrial metabolism and lipid recycling with ExT. **(A)** Top significantly enriched (FDR<0.05) MitoCarta terms from the proteomics sedentary male vs. sedentary female FGSEA results. Points are scaled according to the number of genes in the leading edge subset (the set of genes from each term that contributed to the enrichment score); the top (most upregulated) genes from each leading edge subset are displayed in descending order on the right. Terms were only significantly enriched in males relative to females, so all points are colored blue. **B)** Heatmap of the top MitoCarta terms that were most significantly enriched in any of the 8 trained vs. sedentary control comparisons from the proteomics FGSEA results. **C)** Percentage of reads from mitochondrial genes calculated from the raw transcriptomics data prior to any filtering (top) and the log2-transformed sample-level values of cardiolipin CL(72:8) (bottom). **D)** Proteins involved in lipid metabolism. Values shown are the group means of the standardized sample-level protein values. **E)** Barplot of the median total TAG concentration (μg/mg tissue) in scWAT samples of male and female rats at each timepoint. The heatmap displays the standardized median concentration (peak area normalized by internal standard) of the top 20 most abundant TAG species. **F)** Scatterplots of TAG chain length vs. log2 fold-change from the timewise differential analysis results separated by sex for the 4- and 8-week-trained vs. sedentary control comparisons. Loess curves are included with 95% confidence bands. The y-axes have been restricted to more clearly show patterns, so not all points are visible. **G)** Heatmap of acylcarnitine species grouped according to their metabolomics WGCNA module. Values shown are the group means of the standardized sample-level values for each species.

Specific analysis of proteins regulating scWAT lipid recycling (re-esterification and lipolysis) revealed sexual dimorphism in long-chain acyl-CoA synthetase (Acsl) isoforms, an enzyme family that regulates metabolic partitioning of fatty acyl-CoAs^55^. Females expressed higher levels of the major WAT isoform Acsl1 than males (Fig. 5D). Abundance of glycerol-3-phosphate acyltransferase (Gpat) isoforms (the rate-limiting enzymes of glycerolipid synthesis) also displayed sexual dimorphism, with Gpat3 higher in females compared to males at all timepoints, and Gpat4 higher in males (Fig. 5D). Gpat3 is responsible for ~80% of the enzyme functional activity in WAT and is expressed during adipogenesis^56,57^; conversely, the male-dominant isoform Gpat4 is more abundant in brown adipose tissue (BAT)^57^. Agpat, an enzyme family involved in glycerolipid and phospholipid synthesis, also displayed sexual dimorphism, with Agpat1 and Agpat5 highest in males and Agpat 2–4 highest in females. Increased levels of Agpat2 in females suggests a lipid-storing phenotype, as this is the most abundant Agpat isoform in WAT, while Agpat1 (predominant in males) has bi-directional enzymatic activity in lipid synthesis^58^. Diglyceride acyltransferases (Dgat) and phosphoenolpyruvate carboxykinases (Pck) are also key re-esterification enzymes^59,60^. Consistent with fat mass preservation, females displayed upregulation of Dgat1 and Pck2 at all timepoints (Fig. 5D). The lipid synthesis protein Acaca (Acc1) was elevated in sedentary male scWAT and increased in females following 8 weeks of training (Fig. 5D).

These changes in lipid esterification enzymes were complemented by evidence of sexual dimorphism in the perilipin (Plin) protein family, regulators of LD surface stability and accessibility for lipolysis, as Plin1 was enriched in females and Plin2 in males (Fig. 5D). Plin1 is predominant on the surface of LD of large adipocytes and, in its inactive state, limits LD accessibility and lipolysis^61^. Plin3, which further limits lipolysis, was also elevated in female scWAT^62^. Conversely, the male-dominant Plin2 is associated with small LD^61^ and permits LD accessibility and lipolysis^63^. Together, this suggests that male scWAT is enriched in proteins favoring hydrolysis of lipid droplets. Interestingly, males consistently displayed upregulation of BAT-associated lipid handling protein isoforms, despite not showing upregulation of proteins indicative of scWAT browning or beiging such as Ucp1. We also examined expression of proteins regulating fatty acid oxidation. Males displayed enrichment in Cpt1a and Slc25a20 (Cact) with ExT, proteins key for transport of acylcarnitines into the mitochondria for beta-oxidation (Fig. 5D). Further, males displayed upregulation in other proteins related to lipid oxidation with prolonged training (4 weeks: Cd36, Acsl1, Acsl5, and Fatp1 (Slc27a1); 8 weeks: Acads, Acadvl). Males also displayed increases in proteins involved in lipid peroxisomal alpha-oxidation (Abcd3, Hacl1), LD tethering to peroxisomes (Spast, Abcd1), and markers of peroxisome biogenesis (Pex11b), further emphasizing the catabolic effects of ExT on the male scWAT proteome.

Finally, we examined changes in scWAT TAG abundance and other metabolomic markers of lipid metabolism with ExT. Consistent with decreased fat mass, plasma leptin, and adipocyte size, males decreased median total scWAT TAG abundance by ~70% after 8 weeks of ExT (Fig. 5E). While females did not show substantial changes in scWAT TAG levels over the same time course (Fig. 5E), they exhibited changes in TAG composition, which is indicative of scWAT TAG remodeling with ExT in the absence of reduced fat mass (Figs. 5F, S7B–C). This was characterized by an increase in short chain (SC) and medium chain (MC) TAG abundance in 4- and 8-week-trained females, with decreases in LC TAG species (>40 carbons) at 8 weeks. Conversely, males displayed a reduction in SC TAGs in the early training response, indicated by a reduction in 30–40 carbon species (Fig. S7B–C); by 8 weeks of ExT, the abundance of SC species returned to baseline levels and LC TAG species decreased, albeit to a lesser extent than in females (Figs. 5F, S7B–C). Specific metabolomics modules further support enhanced lipid mobilization and catabolism in males subjected to ExT, leading to depletion of fat stores, versus lipid recycling to preserve fat mass in females. Acyl-CoA and acylcarnitine species measured in this study report on the metabolism of fatty acids. Clear sexual dimorphism was apparent in the acylcarnitine and acyl-CoA profiles with ExT (Figs. 5G, S4A). Females had higher levels of nearly all species of MC and LC (>8 carbons) acylcarnitines at all training timepoints, whereas males had higher levels of SC (<8 carbons) acylcarnitines with the exception of C2 (acetyl) (Fig. 5G). Females also displayed a robust increase in MC and LC acylcarnitines at 1 week, together with overall increased abundance of these species it is suggestive of a reduced capacity for fatty acid oxidation in females, consistent with reduced levels of oxidative proteins. Acyl-CoA levels were also decreased in 4-week-trained males (Fig. S4A). These profiles are consistent with activation of fatty acid oxidation to consume long-chain acylcarnitines and activation of acyl-CoAs to produce SC acylcarnitines in males and preservation of LC species in females, possibly for recycling to TAGs for storage.

## Discussion

We leveraged comprehensive multi-omics datasets generated through the MoTrPAC study of adult rats that began exercise training at 6 months of age to gain deeper insight into scWAT adaptations to progressive endurance training and investigate differences in the response of this tissue between males and females. Multi-omic integration highlighted sexually conserved and sexually dimorphic ExT responses in addition to baseline sexual dimorphism in the scWAT of sedentary rats. We found that male scWAT was enriched in markers of aerobic metabolism and lipid utilization at baseline and with training, contributing to a training-induced decrease in fat stores. Females displayed modulation of markers of scWAT lipid catabolism relative to males and maintenance of fat mass with ExT which may be, in part, due to a compensatory upregulation of mTOR signaling and adipogenesis. WGCNA revealed that sexual dimorphism in the scWAT molecular responses to ExT correlated with sexually distinct phenotypic responses, including changes in key regulatory hormones, metabolites, and cytokines. We also identified differences in lipid-regulatory protein isoforms across sexes, with males displaying an increased abundance of isoforms that are more prevalent in BAT. Together, our findings describe sexual dimorphism in the scWAT progressive response to ExT, providing new insights into factors linking scWAT biology to the risk and pathogenesis of cardiometabolic diseases, especially those that are distinctly impacted by sex.

Although WAT is one of the most sexually dimorphic tissues^22,23^, few studies have examined differential adaptability of scWAT to endurance exercise between sexes in an integrative, multi-omic manner^11,17,18,24–26^. A novel feature of this study is examination of the progressive scWAT ExT response in adult rats, as previous studies have typically evaluated training adaptations at a single time point in young animals. Addressing this gap, our work identified substantial sex-based differences in temporal transcriptional, proteomic, and phosphoproteomic signatures in scWAT in response to ExT. For example, at the transcriptional level, males displayed the highest number of differential features at 2 weeks (529 differential features) of ExT, whilst in females this occurred at 8 weeks (193 differential features); in fact, when compared to females, males had ~66% more differentially regulated transcripts throughout training (788 versus 474). At the proteomic level, males had ~153% more differentially regulated proteins compared to females (1454 versus 574), with the proteomic response peaking at 2 weeks of ExT in females (221 features), and progressively increasing at 4 and 8 weeks in trained males (571 and 654 features, respectively); notably, similar temporal dynamics were observed in the phosphoproteomic signature. Despite this overt sexual dimorphism in response to exercise at the gene and protein level, the metabolomic temporal response to training appeared to be the most consistent between the sexes, with both males and females displaying the greatest number of differential features at 8 (females: 139, males: 211) and 2 weeks (females: 135, males: 161). A particularly unique aspect of our work is cataloging the scWAT metabolomic ExT response and cataloging changes in RefMet subclasses^39^. While this is the first study to define the temporal multi-omic exercise response in males and females, the profound molecular impact of exercise agrees with previous literature demonstrating a strong scWAT ExT response^3,11,12,17,24–26,64,65^.

Importantly, this work provides a multi-omic view of Gene Ontology terms, facilitating broad insight into molecular regulators of the progressive training response. To this point, FGSEA highlighted early (1 week) enrichment of transcriptional and translational regulatory terms in females; the 1-week response in males was enriched for GO terms related to tissue remodeling, including transcripts indicative of immune cell activation and angiogenesis. Both sexes share early increases in PC and O-PC species and terms related to adaptive immune cell activation. Over-representation analysis of WGCNA modules gave results similar to those from FGSEA, indicative of an early adaptive immune response in females and males (T4 ME), which correlated with the plasmalogen-containing M4 and M5 MEs. Plasmalogens are a major phospholipid class of cellular membranes, affecting cellular structural integrity, lipid raft formation, and protein scaffolding, and thus play an important role in transducing cellular signals and adipose remodeling. In lean adipose tissue, immune cells have recently been found to have positive tissue regulatory roles on adipocyte metabolism, lipid utilization and flux^66–69^, where depletion of regulatory T cell subsets contribute to insulin resistance^69^. Examining the role of plasmalogens and immune cell subsets in regulating remodeling of the scWAT microenvironment is therefore ripe for investigation.

The impact of sex on adipose metabolism is likely multifactorial. While sex hormones clearly have an effect on adipogenesis, lipolysis, and re-esterification^19,20,28,70^, other factors such as genetic imprinting and embryonically established tissue-resident cellular compositions likely contribute to sex differences in systemic metabolism^19,23,71^. In the genetically diverse hybrid mouse diversity panel cohort, *ex vivo* WAT mitochondrial function differs based on genetic background in gonadectomized mice in a sex-dependent manner^72^. This emphasizes that gene by sex hormone interactions imprint metabolic phenotypes through undefined mechanisms. In our study of inbred F344 rats, females consistently displayed upregulation of transcripts and proteins related to mRNA processing and ribosome biogenesis that increased with endurance exercise training. Ribosome biogenesis is a tightly regulated process controlling cellular growth in an amino-acid- and MTOR-dependent manner^73^. Depletion of proteins mediating ribosome biogenesis (e.g, Ptfr/Cav1) hinders adipocyte adaptability to changes in nutrient demand^74^; thus, female-specific enrichment of ribosomal gene sets could be an evolutionarily conserved mechanism to sustain adipose allogeneic compatibility with reproductive fitness. Conversely, our study highlights that scWAT in males displays greater tissue plasticity with ExT relative to females, as indicated by the number of differentially regulated features and decreased body fat and scWAT adipocyte size, likely due to an increased need for lipid mobilization in males. Interestingly, the strongest associations between adipocyte size and multi-omic measures were with MEs of the M3 (PC) and M6 (amino acid) WGCNA modules, which were associated with smaller adipocytes and body fat loss. PC has been shown to promote adipocyte lipolysis, inflammation, and apoptosis^75^; however, the role of ExT on PC species remodeling is inconsistent^16,76^. Enrichment of amino acids and MitoCarta terms related to amino acid metabolism in both sexes (1, 2, and 8 weeks in females and 8 weeks in males), may be indicative of switching to an alternate fuel source to spare lipid catabolism with ExT, driving correlation between these measures.

We leveraged phosphoproteomics data to highlight probable molecular transducers of the sexually dimorphic scWAT response to training. With ExT, males displayed enrichment of AMPK activity and females displayed enrichment of AKT and mTOR kinase activity. Adrenaline-mediated AMPKɑ1 signaling promotes lipolysis via ACC phosphorylation ^77^, highlighting a candidate molecular transducer of enhanced lipolytic signaling in male scWAT with ExT. Overall enrichment of AKT and mTOR kinase activity in female scWAT suggests a female-specific mechanism to maintain scWAT adipogenesis and preserve scWAT fat stores in response to ExT^50^. Supporting these observations, mTORC1 signaling promotes scWAT lipid storing capacity and prevents ectopic lipid deposition^78^. Interestingly, ablation of mTOR and LKB prevents diet-induced obesity, yet results in insulin resistance^79^; differences in adipocyte mTOR signaling may therefore regulate sexual dimorphism in adipocyte lipid utilization, impacting differences in insulin sensitivity observed between sexes^19,21,80^. Further supporting enriched anabolic signaling in female scWAT with ExT was the consistent downregulation of the insulin signaling repressor, Grb14, at all training timepoints. Estradiol inhibits insulin-induced Grb14 expression^81^, highlighting a potential mechanism by which female sex hormones promote insulin-mediated anabolic signaling to favor lipogenesis. The lipid-sparing phenotype is additionally evident as the scWAT of sedentary female rats displayed elevated levels of the re-esterification enzymes PCK2 and DGAT1, as well as increased TAG abundance. MitoCarta analyses did highlight enrichment of scWAT OXPHOS proteins in females with training, albeit this response was attenuated and delayed in females relative to males. Despite maintenance of both fat mass and scWAT adipocyte size, female scWAT displayed TAG species remodeling, likely indicative of lipid utilization and subsequent re-esterification. This is supported by WGCNA, which localized transcripts related to neutral lipid biosynthetic processes to the T5 module, the ME of which was highly correlated with change in fat mass and the M1 LC TAG-localizing metabolomics ME. Overall, our integrative multi-omic approach highlights that increased anabolic signaling, likely through the insulin–mTOR axis, maintains fat mass in female rats with ExT.

Our work provides a critical foundation for understanding the coordinated molecular responses to consecutive acute-bout (i.e., chronic) endurance exercise, as these rats were intentionally studied 48 hours following the last bout of exercise. Deciphering acute responses to single bouts of exercise in scWAT, and translation of these findings to outbred rodent and human models are key next steps that can leverage and build upon this work. Our findings highlight that, while largely dimorphic in their training responses, both male and female scWAT displayed metabolically-favorable adaptations to exercise training. This is translationally relevant due to documented sexual dimorphism in fat loss^82^ and fat oxidation^83^ with ExT in humans. While ExT did not lead to fat loss in females, it did prevent excess fat accumulation, as observed in sedentary females. Males significantly reduced fat mass with training through robust increases in lipid utilization which reduces the cardiometabolic risks associated with greater adiposity. Importantly, our study focused on a single subcutaneous fat depot, and other fat depots may respond differently to chronic training^6,20,84,85^. Furthermore, our study design did not control for or measure food intake, so we cannot determine whether the sexual dimorphism in the observed training response was affected by changes in this variable.

In summary, our study is the first to characterize sexually distinct temporal dynamics in scWAT to ExT utilizing an integrative multi-omics approach. Our data highlights early sexually conserved scWAT adaptations, despite sexual dimorphism in sedentary and prolonged training states. Baseline differences in scWAT lipid-regulatory protein isoforms appear to dominate the temporal response to training, with female scWAT displaying increased abundance of proteins related to lipid re-esterification and males displaying upregulation of lipid catabolism and OXPHOS proteins. While we did not observe changes in -omes indicative of ‘browning’ or ‘beiging’, male scWAT displayed increased abundance of lipid droplet-regulatory protein isoforms that typically display higher expression in BAT, albeit expression of ROS-regulatory signaling molecules (e.g., Sod1) were also increased. We hope that our study provides an opportunity for hypothesis-driven investigation of causal links between -omics clusters and phenotypic responses to training, including the impact of sex on such pathways. Identifying sex-specific and sex-conserved scWAT responses to training creates a framework for understanding scWAT cellular regulation, and re-emphasizes the need to consider sexual dimorphism in scWAT to optimize precision-based interventions.

## Methods

### Animals

Adult male and female Fischer 344 (F344) inbred rats were obtained from the National Institute on Aging (NIA) rodent colony. Rats were housed two per cage (146.4 in^2^ of floor space) in ventilated racks (Thoren Maxi-Miser IVC Caging System) on Tekland 7093 Shredded Aspen bedding and fed the LabDiet 5L79 pelleted diet, which are the standard bedding and diet used at the NIA rodent colony. The animal housing room was monitored daily and maintained at a temperature of 68–77°F and humidity of 25–55%. Upon arrival, rats were adapted to a reverse dark–light cycle with lights off at 9 am and lights on at 9 pm for a minimum of 10 days. Red lights were used during the dark cycle to provide adequate lighting for routine housing tasks, rodent handling, and training. All animal procedures were approved by the Institutional Animal Care and Use Committee at the University of Iowa.

### Exercise Training

Exercise training (Panlab 5-lane rat treadmill, Model LE8710RTS, Harvard Instruments) began at 6 months of age and lasted for a duration of 1, 2, 4, or 8 weeks. Rats were exercised on the treadmill 5 days per week using a progressive training protocol designed to exercise the rats at approximately 70% of VO_2_max.

### Body Composition

Body composition was determined for all rats 13 days prior to the start of training using the minispec LF90II Body Composition Rat and Mice Analyzer (Bruker). Post-training body composition was determined for rats in the 4- and 8-week training groups 5 days prior to tissue harvesting.

### Maximal Oxygen Consumption (VO_2_max)

VO_2_max was determined prior to the onset of training in all rats and during the last week of training for the 4- and 8-week exercise groups. Rats were acclimated to a single-lane enclosed treadmill (Columbus Instruments Metabolic Modular Treadmill) 2 days prior to testing. On the day of testing, the rat was placed in the treadmill and testing began once oxygen consumption stabilized. VO_2_, VCO_2_, and respiratory exchange ratio (RER) were recorded at 5-second intervals while in the treadmill; blood lactate levels were recorded at 2-minute intervals. Testing began with a 15-minute warmup at a speed of 9 m/min and 0° incline. Following this, the incline was increased to 10° and treadmill speed was increased by 1.8 m/min every 2 minutes^86^. Post-training males and females remained at speeds of 23.4 and 27 m/min, respectively, for a total of 4 minutes—the first 2 minutes were spent at an incline of 10° and the last 2 at an incline of 15°, which was then maintained for the duration of the testing period. Criteria for reaching VO_2_max was a plateau in oxygen uptake despite increased workload, RER above 1.05, and a non-hemolyzed blood lactate concentration ≥6 mmol/L.

### Statistical Analyses of Phenotypic Measures

Two-sided paired t-tests were performed on each of the phenotypic measures (VO_2_max, body weight, fat mass, etc.) to test for post - pre differences in the sedentary, 4-week, and 8-week-trained groups by sex. The Holm method^87^ was applied to each set of 3 comparisons by sex to control the family-wise error rate.

### Tissue Collection

Tissues were collected from all rats 48 hours following the last training session. On the day of collection, food was removed at 8:30 am, three hours prior to the start of dissections which occurred between 11:30 am and 2:30 pm. Rats were sedated with inhaled isoflurane (1–2%), during which blood was drawn via cardiac puncture, and subcutaneous white fat (inguinal fat depot from right side) was removed and immediately frozen in liquid nitrogen, placed in cryovials, and stored at −80°C. Removal of the heart resulted in death.

### Measurement and Statistical Analyses of Clinical Analytes

We measured a set of nine common clinical analytes in plasma: glucose, lactate, glycerol, total ketones, non-esterified fatty acids (NEFA), glucagon, insulin, leptin, and corticosterone. The first five were measured using a Beckman DxC 600 clinical analyzer with reagents from Beckman (Brea, CA) and Fujifilm Wako (Osaka, Japan; total ketones and NEFA). The others were measured in immunoassays using commercial kits from Meso Scale Discovery (Rockville, MD) and Alpco (Salem, NH; corticosterone). Additionally, we calculated the molar ratio of insulin to glucagon and the Homeostatic Model Assessment for Insulin Resistance (HOMA-IR) by dividing the product of insulin (μIU/mL) and glucose (mg/dL) by 405. For each set of measures, we log-transformed the values and fit a weighted least squares regression model with the reciprocal variances of each sex and timepoint group as weights. Then, each of the trained timepoints were compared to their sex-matched sedentary controls using the Dunnett multiple comparison procedure if timepoint was included as a main effect; if timepoint was not a significant predictor, we instead tested for differences between the sexes. If potential outliers were detected, two sets of results were generated: one with and one without those values. Tests were only performed on the subset of samples selected for multi-omics analyses (n=6 per group). Clinical analyte measures and results of statistical analyses are provided in Supplementary Table 1.

### WAT Histology

scWAT tissue samples were collected from the contralateral side of rats used for multi-omics analyses, embedded in OCT, and flash frozen for histological analyses. Tissue samples were sectioned with a thickness of 15 μm in a cryostat, stained with hematoxylin and eosin (H&E), and visualized with bright field microscopy. A total of 10 fields of view were imaged per sample using automated CellProfiler software to quantify adipocyte cell size^88^. Adipocytes were binned in 5 μm intervals according to their diameter. A log-link negative binomial regression model that included sex, timepoint, diameter bin (ordered factor, 10 levels), and their interactions as predictors was used to model the expected rate of adipocytes (offset = log(total adipocytes per sex and timepoint group)). Within each bin, we compared the 4- and 8-week-trained timepoints to their sex-matched sedentary controls using the Dunnett multiple comparison procedure. The ratios of the expected values are shown in Figure S1D if the adjusted p-values were below 0.05. Measures of adipocyte diameter, area, and volume, as well as results of the statistical analysis are provided in Supplementary Table 1.

### Multi-omic Data Generation and Processing

Full details of the methods used for sample processing, data collection, processing, normalization, and batch correction for transcriptomics, proteomics, phosphoproteomics and metabolomics/lipidomics platforms are described elsewhere^22^, and summarized in Supplementary Text.

### Removal of Redundant Metabolites

A total of 1286 metabolites were measured in scWAT across 13 targeted and untargeted metabolomics platforms and 6 analysis sites, as outlined in Methods Table 4 of MoTrPAC Study Group^22^, with 353 metabolites that were measured in multiple platforms. If metabolites were measured in both targeted and untargeted assays, only those from the targeted assays were retained. The remaining redundant features (detected in multiple untargeted assays) were visually inspected and selection was made case-by-case based on metabolite polarity, solubility, charge, and, therefore, their suitable sample preparation (e.g., extraction solvent) and chromatography method (e.g., column). This resulted in a total of 1063 unique metabolites for analysis. Complete details are included in the Supplementary Note.

### Differential Analysis

RNA-Seq data was analyzed using the *edgeR^89^* (v3.40.1) and *limma^90^* (v3.54.0) Bioconductor/R^91,92^ packages, following the workflow described by Law *et al*.^93^. Briefly, low-abundance transcripts were removed from the raw count data with *edgeR∷filterByExpr* to more accurately estimate the mean–variance relationship in the data. Then, a multidimensional scaling (MDS) plot was generated from the log_2_ TMM-normalized counts per million reads to explore the average log_2_ fold-change between samples (Fig. S8A). Two samples (90423017005 and 90410017005) were identified as outliers and removed^22^. Furthermore, higher variability was observed in females relative to males, and biological replicates (e.g., all 4-week-trained males) appeared to cluster poorly. To account for this, *limma∷voomWithQualityWeights* was chosen to simultaneously combine observation-level weights (predicted from the mean–variance trend in Fig. S8B) with sample-specific quality weights. These weights are incorporated into the linear model to down-weight low abundance observations and all observations from more variable samples, increasing power to detect differences^94,95^.

Proteomics, phosphoproteomics, and metabolomics data were also analyzed using *limma*, and differences in sample variability were again apparent from their respective MDS plots (Fig. S8C–E). As such, sample-specific quality weights were calculated with *limma∷arrayWeights* (*method=“genebygene”*) and incorporated into the linear models^96^. Additionally, RNA integrity number, median 5’-3’ bias, percent of reads mapping to globin, and percent of PCR duplicates as quantified with unique molecular identifiers were included as covariates in the RNA-Seq model after they had been mean-imputed and standardized^22^. For each -ome, we set up a no-intercept model containing the experimental group (each combination of sex and timepoint; factor with 10 levels) and any covariates.

Following linear modeling with *limma∷lmFit*, contrasts were constructed with *limma∷contrasts.fit* to test sex-specific training responses (e.g, 1W males vs SED males; Supplementary Table 4), baseline sexual dimorphism (SED males vs SED females; Supplementary Table 2), and sexually dimorphic training responses (i.e., the sex by training interaction effect). Then, robust Empirical Bayes moderation was carried out with *limma∷eBayes* to squeeze the residual variances toward a common value (RNA-Seq) or a global trend (proteomics and phosphoproteomics; Fig. S8F–G)^97^. For metabolomics, the Empirical Bayes moderation was performed separately for each of the 13 platforms with *robust=trend=TRUE* (both *FALSE* if a platform measured fewer than 10 metabolites).

To control the false discovery rate (FDR), p-values were adjusted across sets of related comparisons by -ome using the Benjamini–Hochberg (BH) procedure^98^. Counts of differential features (FDR < 0.05) from the timewise comparisons are shown in the UpSet plots (Fig. 3A–B)^99^, which were generated with the *ComplexHeatmap* R/Bioconductor package (v2.12.0)^100^.

### Fast Gene Set Enrichment Analysis

Gene Set Enrichment Analysis (GSEA) is a rank-based approach that determines whether any *a priori* defined gene sets—such as genes involved in the same biological process or participating in the same pathway—display concordant behavior^101,102^. FGSEA (specifically, FGSEA*-multi-level*) is a fast implementation of preranked/gene permutation GSEA based on an adaptive multi-level split Monte Carlo scheme for the estimation of arbitrarily small p-values^38^. It requires gene-level statistics, known as ranking metric values, and a collection of gene sets. We chose the signed -log_10_-transformed p-value (calculated from the differential analysis results for each contrast) as the ranking metric, where the sign indicates the direction of the log_2_ fold-change; while similar to the t-statistic, this metric more clearly separates extreme values from those near zero.

For each contrast, ranking metric values were calculated at the level of individual features (i.e., proteins, transcripts, metabolites). Proteomics and transcriptomics ranking metrics were aggregated to the Entrez gene ID level by taking the arithmetic mean, and any features that did not map to an Entrez gene were discarded prior to analysis.

For proteomics and transcriptomics, gene sets from three C5:GO subcollections—Biological Process (BP), Molecular Function (MF), and Cellular Component (CC)^103,104^—of the Molecular Signatures Database^105^ (MSigDB v7.5.1) as provided by the *msigdbr^106^* R package were considered for testing. These collections were chosen, in part, because they have been filtered to reduce inter-set redundancy using a method similar to the one described by Liberzon *et al*.^105^; the exact method is outlined in sections 3.2 and 3.7 of the v7.0 MSigDB Release Notes on the Broad Institute’s website (https://software.broadinstitute.org/cancer/software/gsea/wiki/index.php/MSigDB_v7.0_Release_Notes). After filtering each set to the genes from a particular -ome, they were required to have retained at least 85% of their original members and to contain at least 15 and no more than 300 genes. The high membership percentage cutoff was imposed due to the high proteome (9964 proteins) and transcriptome (16404 transcripts) coverage that was observed for this tissue. Set size was restricted because smaller sets are less reliable while larger sets are more difficult to interpret.

For proteomics only, we additionally tested gene sets from the MitoCarta3.0 database (file “Human.MitoCarta3.0.xls”) (Fig. 5C–D). Human gene symbols were remapped to their rat orthologs (Entrez gene IDs) prior to analysis. The same 85% membership filter was applied, though terms were only required to contain at least 5 and no more than 300 genes due to their already small sizes; this resulted in 63 sets for testing.

Finally, the FGSEA procedure was applied to the metabolomics data by grouping metabolites according to RefMet chemical subclasses (e.g., acylcarnitines and triacylglycerols)—provided by the Metabolomics Workbench RefMet database (https://www.metabolomicsworkbench.org)^39^. Since these subclasses are disjoint and homogenous, they were not filtered based on a membership percentage, though each subclass was required to contain at least 10 metabolites.

FGSEA was performed with *fgsea∷fgseaMultiLevel* (v1.24.0)^38^ in R. A total of 10,000 permutations were used for the preliminary estimation of enrichment p-values and to calculate normalized enrichment scores (NES), and p-values were adjusted across groups of related contrasts within each -ome and feature set collection (e.g., across all proteomics GO:MF results) using the BH procedure^98^. The enrichment heatmaps (Figs. 3, S3, S5), created with the *ComplexHeatmap* (v2.14.0)^100^ R/Bioconductor package, display the terms with the highest -log_10_-transformed adjusted p-value in any contrast. The circles are scaled such that the lowest adjusted p-value in each row is of maximum area; this is to prevent one or more feature sets with extremely low adjusted p-values from dominating the heatmaps. NES in [−1, 1] are essentially noise^38,102^, so those circles are not shown. Results are provided in Supplementary Tables 3 and 5.

### Kinase–Substrate Enrichment Analysis

To infer kinase activity from our phosphoproteomics data, we leveraged the curated kinase–substrate information provided by PhosphoSitePlus^45^ (PSP; v6.6.0.4). As data is significantly more comprehensive for humans than for rats, we first remapped the 30,304 quantified phosphorylation sites on rat proteins to human orthologs using the data generated by the MoTrPAC Study Group^22^ (Methods: Mapping PTMs from rat to human proteins). This resulted in 19,137 unique phosphorylation sites on human proteins. FGSEA^38^ was then performed for each of the phosphoproteomics contrasts with *fgsea∷fgseaMultiLevel* in R using the single-site-level signed -log_10_-transformed p-value as the ranking metric. Kinase sets were constructed from the PSP kinase–substrate dataset (Kinase_Substrate_Dataset.xlsx, accessed 2022-06-05 from https://www.phosphosite.org/staticDownloads) by grouping sites phosphorylated by the same kinase. Only those kinases with at least 3 annotated substrate sites (excluding instances of autophosphorylation) were tested for enrichment. As with previous FGSEA, a total of 10,000 permutations were used for the preliminary estimation of enrichment p-values and NES, and p-values were adjusted across all kinases with the BH procedure^98^.

This approach is similar to that of Ochoa *et al*.^107^, though with several notable improvements owing to the more sophisticated FGSEA algorithm and up-to-date kinase–substrate relationship data. Namely, enrichment p-values are now based on normalized enrichment scores, which account for differences in feature set size, and they are no longer bounded by 0.001 (requiring at least 1/1000 permutation scores to be more extreme). Furthermore, we are able to impose limits on set size to exclude smaller, less reliable sets. This analysis is still limited by the coverage of PSP, however, as the kinases of only 1,074 of the 19,173 phosphosites were known, and only 121/379 kinases passed the size filter for testing. Results are provided in Supplementary Tables 3 and 5.

### Gene Set Network Diagrams

We utilized Cytoscape (v3.9.1)^108,109^ to generate a network diagram (Fig. 2D) of the top significantly enriched biological processes (BH-adjusted p-value < 0.05) from the sedentary male versus sedentary female proteomics FGSEA results. The combined Jaccard and overlap coefficient^110^ was used to calculate edge weights, and only edge weights ≥0.25 were retained. Clusters of nodes were manually assigned summary descriptors, and only the top enriched groups are shown.

### Weighted Gene Co-expression Network Analysis

Weighted gene co-expression network analysis (WGCNA) was carried out with the *WGCNA* R package (v1.71)^111,112^ to identify non-overlapping groups of related proteins, metabolites, and transcripts. Signed adjacency matrices were constructed from the pairwise protein, metabolite, or transcript biweight midcorrelations^113^ using a soft-thresholding power of 25 for transcriptomics and 12 for both proteomics and metabolomics/lipidomics. Additionally, transcriptomics count data was converted to log_2_ TMM-normalized counts per million reads with *edgeR∷cpm*, per the workflow described by Law *et al*.^93^ (Differential Analysis section). No other processing was performed prior to WGCNA.

Average linkage hierarchical clustering was performed on the dissimilarity matrices—computed from unsigned topological overlap matrices—and *dynamicTreeCut∷cutreeDynamic* (v1.63.1)^114^ was used to define the protein, metabolite/lipid, and transcript modules with non-default parameters *cutHeight=0.99*, *deepSplit=2*, *minClusterSize=30*, and *pamRespectsDendro=FALSE*. A total of 6 metabolites and 2587 transcripts were not co-expressed (assigned to “grey” modules) and subsequently discarded. The remaining 1057 metabolites/lipids were assigned to 7 modules of size 30 to 415, the 9964 quantified proteins to 7 modules of size 165 to 3984, and the 13817 transcripts to 14 modules of size 33 to 4683. The modules are arranged in descending order by size and labeled with the first letter of their respective -ome (M, T, or P). Module eigenfeatures (MEs; generalization of module eigengenes for non gene-centric data)—the principal eigenvector of each module^113^—were extracted and Pearson correlations were calculated between metabolomics/lipidomics and proteomics MEs, between metabolomics/lipidomics and transcriptomics MEs (Fig. 4B), and between each ME and select sample measures (Fig. 4A). The significance of each correlation was assessed with *WGCNA∷corPvalueStudent* and adjusted within each correlation matrix using the BH procedure^98^. The correlation heatmaps were created with the *ComplexHeatmap* R/Bioconductor package (v2.14.0)^100^.

### WGCNA Module Over-representation Analysis

In order to characterize the WGCNA modules, over-representation analysis (ORA)^53^ was performed in R with *fgsea∷fora^38^* on the same feature sets that were used for FGSEA. All Entrez or RefMet IDs from the appropriate WGCNA results (excluding those from the “grey” modules) were set as the background for these hypergeometric tests. Since the resulting p-value histograms exhibited peaks near both 0 and 1 (indicating under-represented sets), and there appeared to be an association between p-value and feature set size, p-values were not adjusted to control the FDR. Instead, an overlap ratio (equation (1)) was calculated by dividing the cardinality of the intersection between each module and feature set by one plus the cardinality of the module-wise maximum intersection. (Adding 1 to the denominator penalizes small maximum intersections.)

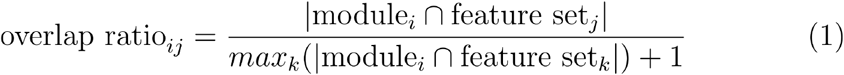

The over-representation p-values were then log_10_-transformed, multiplied by their associated overlap ratio, and back-transformed. This removed the association between set size and p-value, and the histograms of these scaled p-values appeared left-skewed. A small scaled p-value indicates that a feature set is over-represented in a particular module and explains it well relative to all terms that were tested. The top over-represented (scaled p-value < 0.05) feature sets by module are shown in Figure 4C–E. Full results are provided in Supplementary Table 6.

### Mitochondrial DNA Quantification

Quantification of mitochondrial DNA (mtDNA) was performed and described by Amar, *et al*.^115^. Briefly, real-time quantitative PCR was performed in duplicate for each of the scWAT samples selected for -omics analysis. The 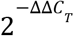 method^116^ was then applied to estimate the relative expression of the mitochondrial D-loop. Since both target (D-loop) and internal control (β-actin) were amplified in the same well, Δ*C*_T_ was calculated as the mean of (*C*_T,D-loop_ - *C*_T,β-actin_) for each sample. Then, ΔΔ*C*_T_ values were obtained by subtracting each Δ*C*_T_ value by the mean Δ*C*_T_ of the sedentary female group (the calibrator). Kruskal–Wallis tests were performed on the 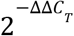 values separately by sex.

## Supporting information

MoTrPAC Study Group Listing

## Abbreviations

ACLY: ATP citrate lyase
Acsl: long-chain acyl CoA synthetase
AMP, ATP: adenosine mono/tri-phosphate
BAT: brown adipose tissue
BCAA: branched-chain amino acid
BH: Benjamini–Hochberg
DAG: diacylglycerol
Dgat: diglyceride acyltransferase
DNL: de novo lipogenesis
ExT: endurance exercise training
FA: fatty acid
FDR: false discovery rate
FGSEA: fast gene set enrichment analysis
GO-(BP, CC, MF): Gene Ontology (biological process; cellular component; molecular function)
Gpat: glycerol-3-phosphate acyltransferase
GSEA: gene set enrichment analysis
H&E: hematoxylin and eosin
HOMA-IR: homeostatic model assessment for insulin resistance
KSEA: kinase–substrate enrichment analysis
LC, MC, SC: long chain, medium chain, short chain
LD: lipid droplet
LPC: lysophosphatidylcholine
MDS: multidimensional scaling
ME: module eigenfeature
METAB: metabolomics/lipidomisc
MoTrPAC: Molecular Transducers of Physical Activity Consortium
MSigDB: Molecular Signatures Database
mtDNA: mitochondrial DNA
NEFA: non-esterified fatty acid
NES: normalized enrichment score
NIA: National Institute on Aging
NMR: nuclear magnetic resonance
O-PC: ether phosphatidylcholine
ORA: over-representation analysis
OXPHOS: oxidative phosphorylation
PC: phosphatidylcholine
PCA: principal component analysis
Pck: phosphoenolpyruvate carboxykinase
PE: phosphatidylethanolamine
PHOSPHO: phosphoproteomics
Plin: perilipin
PROT: proteomics
PSP: PhosphoSitePlus
qPCR: quantitative polymerase chain reaction
RER: respiratory exchange ratio
scWAT: subcutaneous white adipose tissue
SED: sedentary control
SM: sphingomyelin
TAG: triacylglycerol
TRNSCRPT: transcriptomics
VO_2_max: maximal oxygen consumption
WAT: white adipose tissue
WGCNA: weighted gene co-expression network analysis

## Data and code availability

Processed data and analysis results are available in the MotrpacRatTraining6moWATData R package (https://github.com/PNNL-Comp-Mass-Spec/MotrpacRatTraining6moWATData). R scripts detailing all processing and analysis steps are contained within the data-raw/ folder of this package. Helper functions used to perform analyses and visualize results are contained within the MotrpacRatTraining6moWAT R package (https://github.com/PNNL-Comp-Mass-Spec/MotrpacRatTraining6moWAT). MoTrPAC data will be publicly available at time of publication via motrpac-data.org/data-access. Data access inquiries should be sent to. Additional resources can be found at https://motrpac.org and https://motrpac-data.org.

## Acknowledgements

**Funding**: The MoTrPAC Study is supported by NIH grants U24OD026629 (Bioinformatics Center), U24DK112349, U24DK112342, U24DK112340, U24DK112341, U24DK112326, U24DK112331, U24DK112348 (Chemical Analysis Sites), U01AR071133, U01AR071130, U01AR071124, U01AR071128, U01AR071150, U01AR071160, U01AR071158 (Clinical Centers), U24AR071113 (Consortium Coordinating Center), U01AG055133, U01AG055137 and U01AG055135 (PASS/Animal Sites). This work was also supported by other funding sources: NHGRI Institutional Training Grant in Genome Science 5T32HG000044 (N.R.G.), National Science Foundation Graduate Research Fellowship Grant No. NSF 1445197 (N.R.G.), National Heart, Lung, and Blood Institute of the National Institute of Health F32 postdoctoral fellowship award F32HL154711 (P.M.J.B.), the Knut and Alice Wallenberg Foundation (M.E.L.), National Science Foundation Major Research Instrumentation (MRI) CHE-1726528 (F.M.F.), National Institute on Aging P30AG044271 and P30AG003319 (N.M.), and National Institute of Diabetes and Digestive and Kidney Diseases Ruth L. Kirchstein National Research Service Award F32-DK126432 (M.V.).

Parts of this work were performed in the Environmental Molecular Science Laboratory, a U.S. Department of Energy national scientific user facility at Pacific Northwest National Laboratory in Richland, Washington.

We would also like to thank Dr. Surendra Dasari, Dr. Karyn Esser, Dr. Andrea Hevener, and Dr. Sarah Lessard for critical internal MoTrPAC review of this manuscript prior to submission.

The views expressed are those of the authors and do not necessarily reflect those of the NIH or the Department of Health and Human Services of the United States.

## Summary Figure

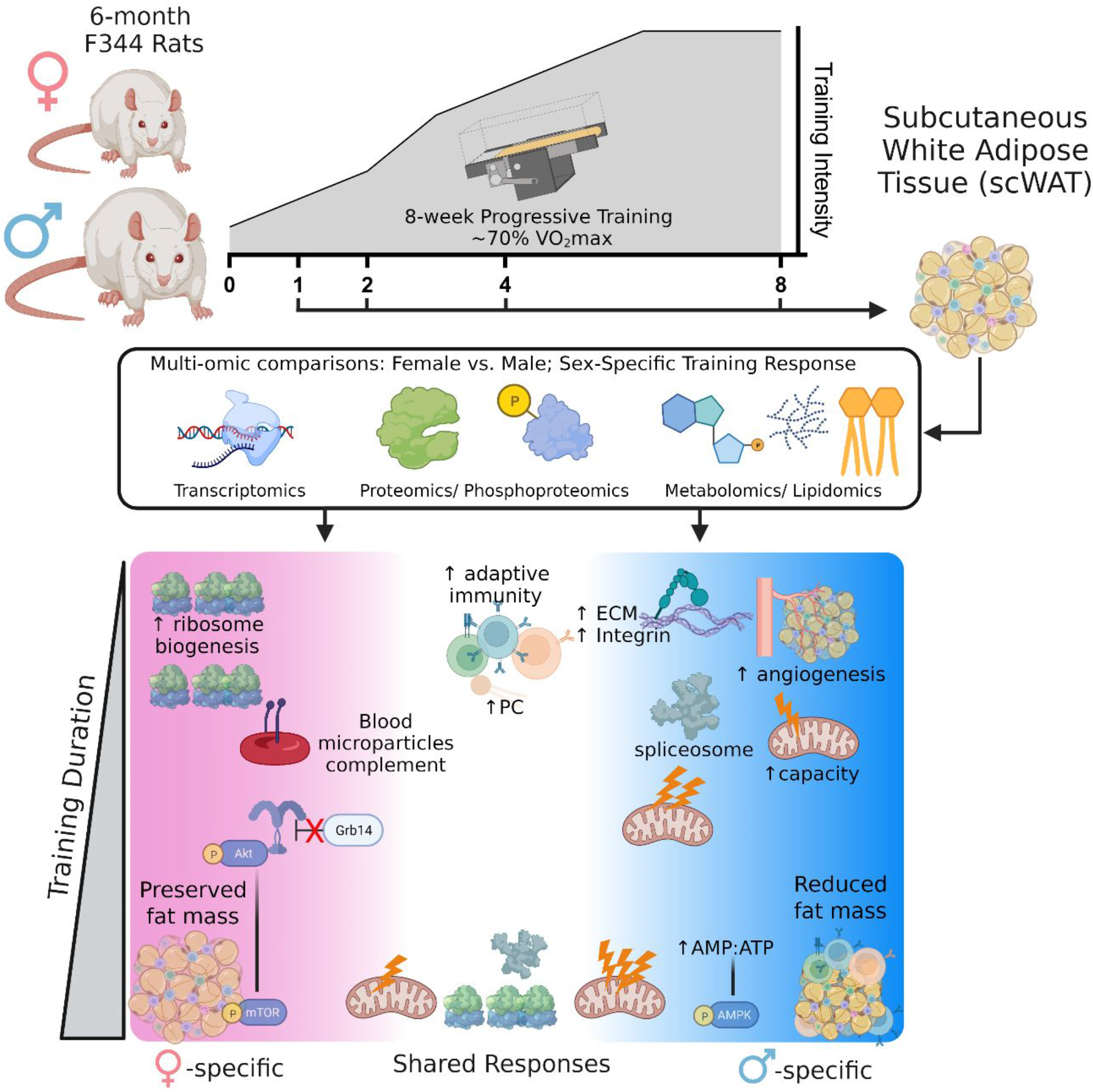

## Supplementary Materials

- **Supplementary Figures**. Supplementary Figures S1–S8 and corresponding figure legends
- **Supplementary Table 1**. Data and statistics for clinical analytes measured in plasma; data and statistics for adipocyte histology
- **Supplementary Table 2**. Complete differential expression analysis (DEA) results for sedentary male vs. sedentary female comparisons in each omics dataset
- **Supplementary Table 3**. Complete fast gene set enrichment analysis (FGSEA) results for sedentary male vs. sedentary female comparisons in each omics dataset
- **Supplementary Table 4**. Complete differential expression analysis (DEA) results for sex-specific exercise training response comparisons in each omics dataset
- **Supplementary Table 5**. Complete fast gene set enrichment analysis (FGSEA) results for sex-specific exercise training response comparisons in each omics dataset
- **Supplementary Table 6**. Complete over-representation analysis (ORA) results for all WGCNA modules from transcriptomics, proteomics and metabolomics datasets
- **Supplementary Text**. Overview of sample preparation, data generation and data processing methods used for multi-omics analysis
- **Supplementary Note**. Details on the selection criteria and process used to remove redundant metabolites identified and quantified on multiple metabolomics platforms
- **MoTrPAC Study Group Listing**

