## Supplementary Figures for "Sexual dimorphism and the multi-omic response to exercise training in rat subcutaneous white adipose tissue"

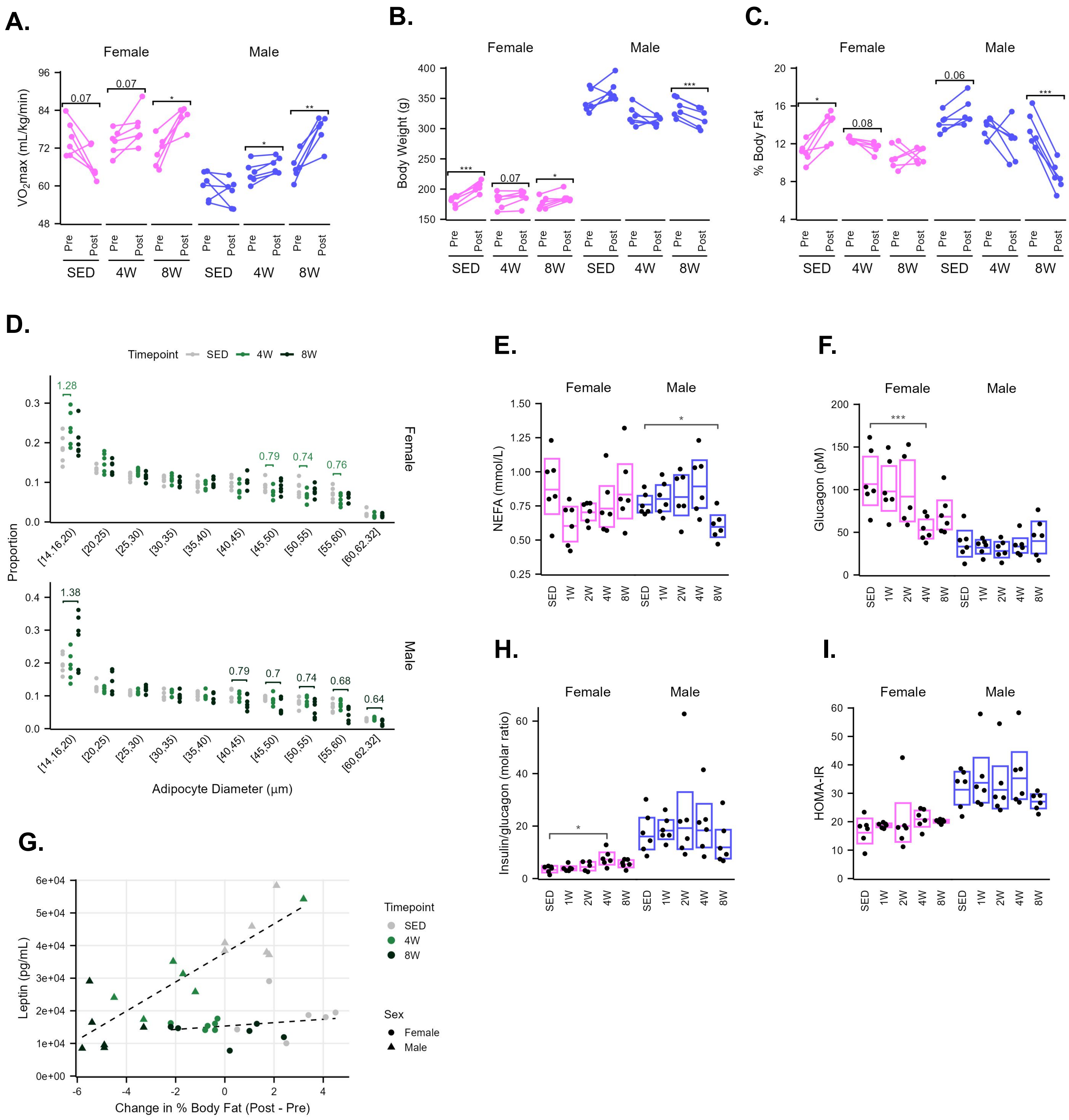

A.
Female
Male
VO2max (mL/kg/min)
96
84
72
60
48
0.07
0.07
*
*
**
Pre
Post
SED
4W
8W
SED
4W
8W
B.
Female
Male
Body Weight (g)
400
350
300
250
200
150
***
0.07
*
***
C.
Female
Male
% Body Fat
20
16
12
8
4
*
0.08
0.06
***
D.
Timepoint
SED
4W
8W
Proportion
1.28
0.79
0.74
0.76
Female
1.38
0.79
0.7
0.74
0.68
0.64
Male
[14,16,20)
[20,25)
[25,30)
[30,35)
[35,40)
[40,45)
[45,50)
[50,55)
[55,60)
[60,62,32]
Adipocyte Diameter (μm)
E.
Female
Male
NEFA (mmol/L)
1.50
1.25
1.00
0.75
0.50
0.25
*
SED
1W
2W
4W
8W
F.
Female
Male
Glucagon (pM)
200
150
100
50
0
***
G.
Leptin (pg/mL)
6e+04
5e+04
4e+04
3e+04
2e+04
1e+04
0e+00
-6
-4
-2
0
2
4
Change in % Body Fat (Post - Pre)
Timepoint
SED
4W
8W
Sex
Female
Male
H.
Female
Male
Insulin/glucagon (molar ratio)
60
40
20
0
*
I.
Female
Male
HOMA-IR
60
50
40
30
20
10
0

**Supplementary Figure S1. Additional phenotypic data.**
**A**) Relative $VO_2$max values normalized to total body mass from the animals selected for multi-omics analysis. **B–C**) Body weight and percent body fat measurements of animals selected for multi-omics analysis as measured by NMR. **D)** Proportions of adipocytes from histological analysis with diameters in the indicated ranges for both female and male animals after 4 or 8 weeks of training, or sedentary controls. For comparisons between trained and sedentary groups where statistical significance was reached, the ratio between the group means (trained/sedentary) is specified above the appropriate bin. **E–F)** Additional measures of plasma clinical analytes: non-esterified fatty acids (NEFA) (**E**) and glucagon (**F**). **G)** Correlation between plasma leptin measurements and changes in percent body fat (post vs. pre-training) in each SED, 4W, and 8W animal selected for -omics analysis. **H–I)** Additional measures of clinical analytes in plasma: molar insulin to glucagon ratio (**H**) and HOMA-IR score (**I**).

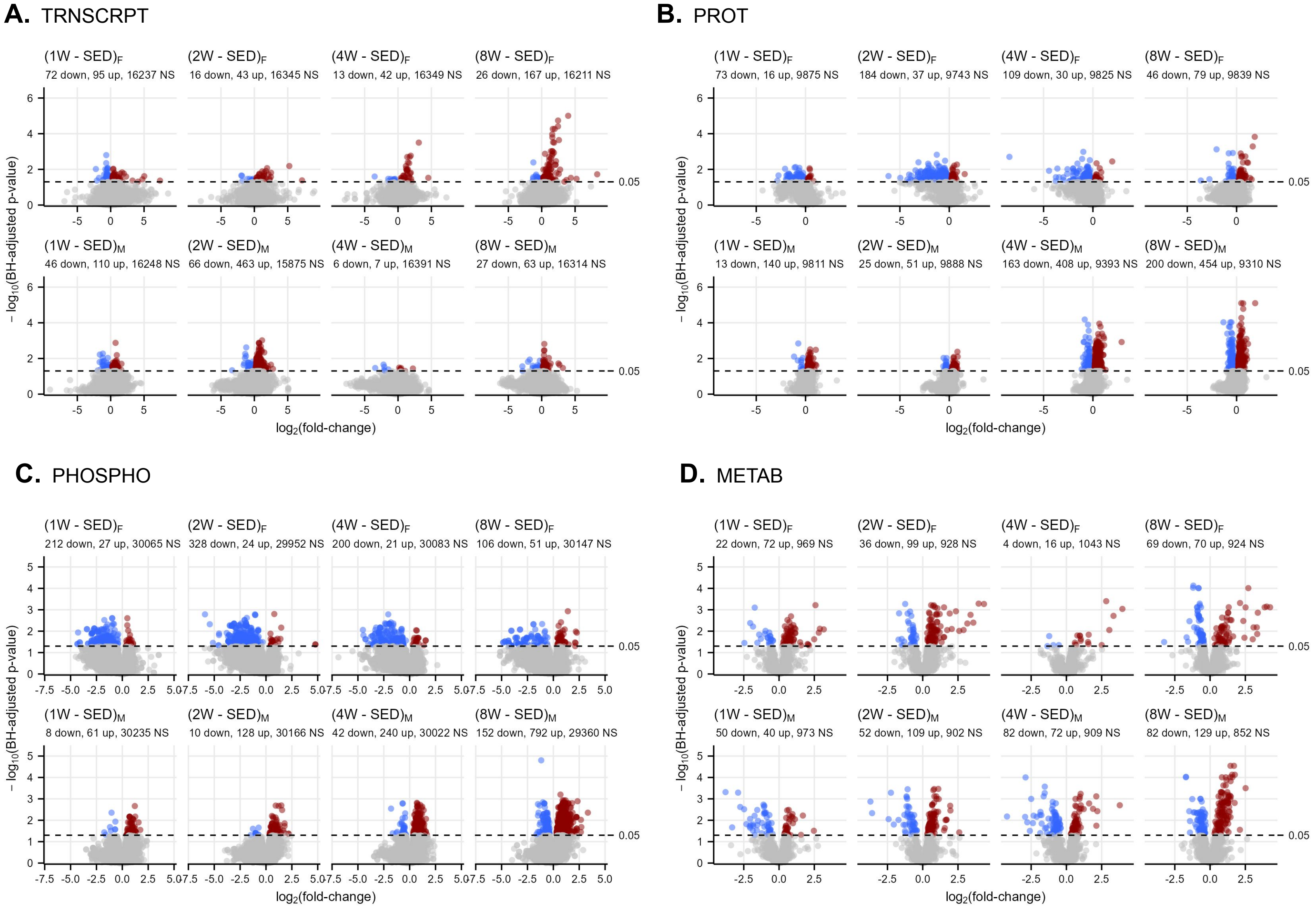
A. TRNSCRPT
(1W - SED)F
72 down, 95 up, 16237 NS
(2W - SED)F
16 down, 43 up, 16345 NS
(4W - SED)F
13 down, 42 up, 16349 NS
(8W - SED)F
26 down, 167 up, 16211 NS
(1W - SED)M
46 down, 110 up, 16248 NS
(2W - SED)M
66 down, 463 up, 15875 NS
(4W - SED)M
6 down, 7 up, 16391 NS
(8W - SED)M
27 down, 63 up, 16314 NS
B. PROT
(1W - SED)F
73 down, 16 up, 9875 NS
(2W - SED)F
184 down, 37 up, 9743 NS
(4W - SED)F
109 down, 30 up, 9825 NS
(8W - SED)F
46 down, 79 up, 9839 NS
(1W - SED)M
13 down, 140 up, 9811 NS
(2W - SED)M
25 down, 51 up, 9888 NS
(4W - SED)M
163 down, 408 up, 9393 NS
(8W - SED)M
200 down, 454 up, 9310 NS
C. PHOSPHO
(1W - SED)F
212 down, 27 up, 30065 NS
(2W - SED)F
328 down, 24 up, 29952 NS
(4W - SED)F
200 down, 21 up, 30083 NS
(8W - SED)F
106 down, 51 up, 30147 NS
(1W - SED)M
8 down, 61 up, 30235 NS
(2W - SED)M
10 down, 128 up, 30166 NS
(4W - SED)M
42 down, 240 up, 30022 NS
(8W - SED)M
152 down, 792 up, 29360 NS
D. METAB
(1W - SED)F
22 down, 72 up, 969 NS
(2W - SED)F
36 down, 99 up, 928 NS
(4W - SED)F
4 down, 16 up, 1043 NS
(8W - SED)F
69 down, 70 up, 924 NS
(1W - SED)M
50 down, 40 up, 973 NS
(2W - SED)M
52 down, 109 up, 902 NS
(4W - SED)M
82 down, 72 up, 909 NS
(8W - SED)M
82 down, 129 up, 852 NS
− log10(BH-adjusted p-value)
log2(fold-change)
0.05

**Supplementary Figure S2. Training response differential expression analysis: volcano plots.**
**A–D**) Volcano plots displaying the magnitude and significance of comparisons of each trained group against sex-matched sedentary controls in transcriptomics (**A**), proteomics (**B**), phosphoproteomics (**C**), and metabolomics (**D**) datasets.

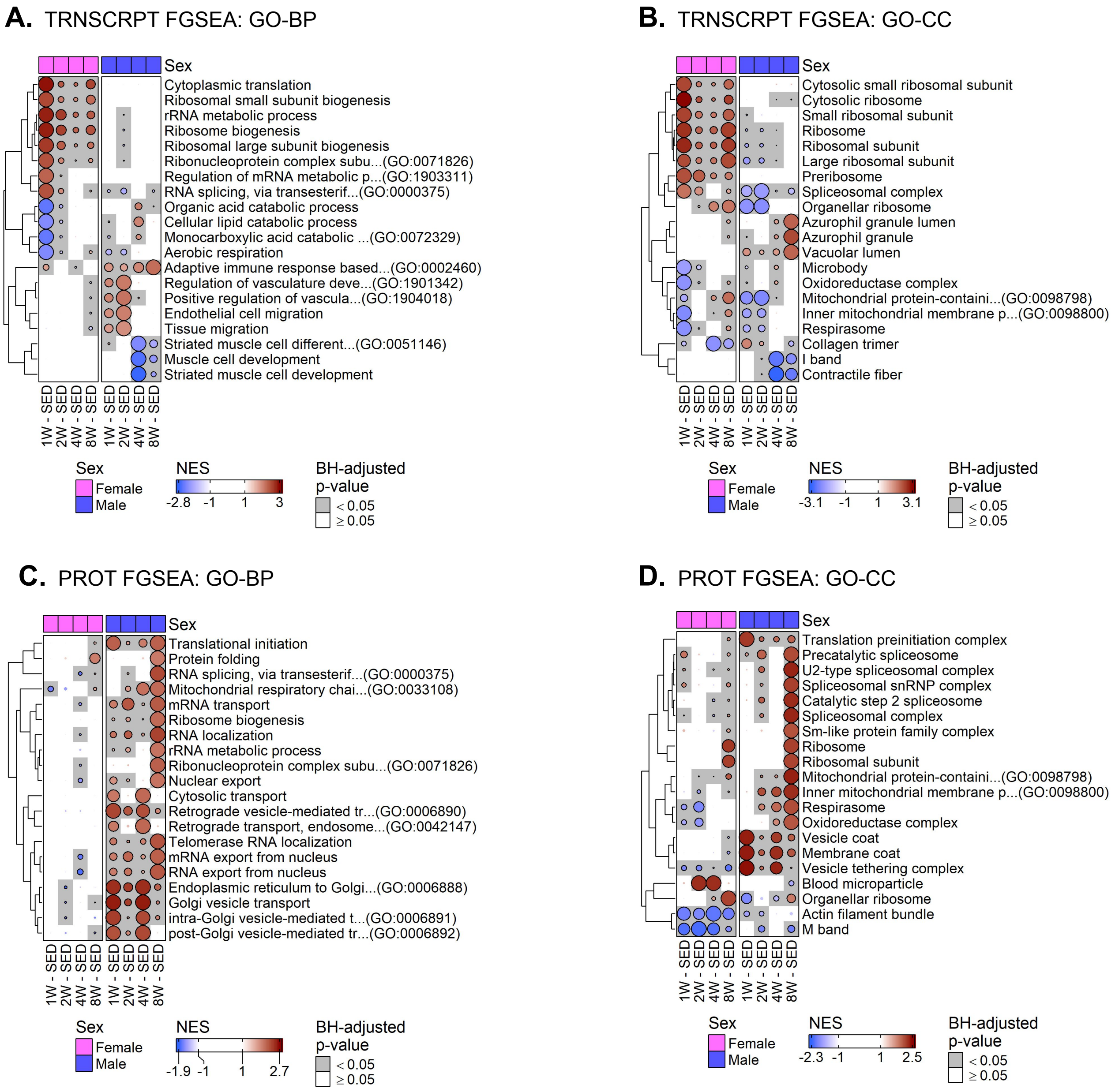
A. TRNSCRPT FGSEA: GO-BP
Sex
Cytoplasmic translation
Ribosomal small subunit biogenesis
rRNA metabolic process
Ribosome biogenesis
Ribosomal large subunit biogenesis
Ribonucleoprotein complex subu...(GO:0071826)
Regulation of mRNA metabolic p...(GO:1903311)
RNA splicing, via transesterif...(GO:0000375)
Organic acid catabolic process
Cellular lipid catabolic process
Monocarboxylic acid catabolic ...(GO:0072329)
Aerobic respiration
Adaptive immune response based...(GO:0002460)
Regulation of vasculature deve...(GO:1901342)
Positive regulation of vascula...(GO:1904018)
Endothelial cell migration
Tissue migration
Striated muscle cell different...(GO:0051146)
Muscle cell development
Striated muscle cell development
1W - SED
2W - SED
4W - SED
8W - SED
Female
Male
NES
-2.8
-1
1
3
BH-adjusted p-value
< 0.05
≥ 0.05
B. TRNSCRPT FGSEA: GO-CC
Sex
Cytosolic small ribosomal subunit
Cytosolic ribosome
Small ribosomal subunit
Ribosome
Ribosomal subunit
Large ribosomal subunit
Preribosome
Spliceosomal complex
Organellar ribosome
Azurophil granule lumen
Azurophil granule
Vacuolar lumen
Microbody
Oxidoreductase complex
Mitochondrial protein-containi...(GO:0098798)
Inner mitochondrial membrane p...(GO:0098800)
Respirasome
Collagen trimer
I band
Contractile fiber
1W - SED
2W - SED
4W - SED
8W - SED
Female
Male
NES
-3.1
-1
1
3.1
BH-adjusted p-value
< 0.05
≥ 0.05
C. PROT FGSEA: GO-BP
Sex
Translational initiation
Protein folding
RNA splicing, via transesterif...(GO:0000375)
Mitochondrial respiratory chai...(GO:0033108)
mRNA transport
Ribosome biogenesis
RNA localization
rRNA metabolic process
Ribonucleoprotein complex subu...(GO:0071826)
Nuclear export
Cytosolic transport
Retrograde vesicle-mediated tr...(GO:0006890)
Retrograde transport, endosome...(GO:0042147)
Telomerase RNA localization
mRNA export from nucleus
RNA export from nucleus
Endoplasmic reticulum to Golgi...(GO:0006888)
Golgi vesicle transport
intra-Golgi vesicle-mediated t...(GO:0006891)
post-Golgi vesicle-mediated tr...(GO:0006892)
1W - SED
2W - SED
4W - SED
8W - SED
Female
Male
NES
-1.9
-1
1
2.7
BH-adjusted p-value
< 0.05
≥ 0.05
D. PROT FGSEA: GO-CC
Sex
Translation preinitiation complex
Precatalytic spliceosome
U2-type spliceosomal complex
Spliceosomal snRNP complex
Catalytic step 2 spliceosome
Spliceosomal complex
Sm-like protein family complex
Ribosome
Ribosomal subunit
Mitochondrial protein-containi...(GO:0098798)
Inner mitochondrial membrane p...(GO:0098800)
Respirasome
Oxidoreductase complex
Vesicle coat
Membrane coat
Vesicle tethering complex
Blood microparticle
Organellar ribosome
Actin filament bundle
M band
1W - SED
2W - SED
4W - SED
8W - SED
Female
Male
NES
-2.3
-1
1
2.5
BH-adjusted p-value
< 0.05
≥ 0.05

**Supplementary Figure S3. Additional FGSEA heatmaps from training response differential expression analysis.**

**A–B)** Top biological process (GO-BP) (**A**) or cellular component (GO-CC) (**B**) terms from the Gene Ontology database that are most significantly enriched in any of the 8 comparisons from transcriptomics FGSEA results. **C–D)** Top GO-BP (**C**) or GO-CC (**D**) terms that are most significantly enriched in any of the 8 comparisons from proteomics FGSEA results.

**A.** scWAT fatty acyl-CoAs

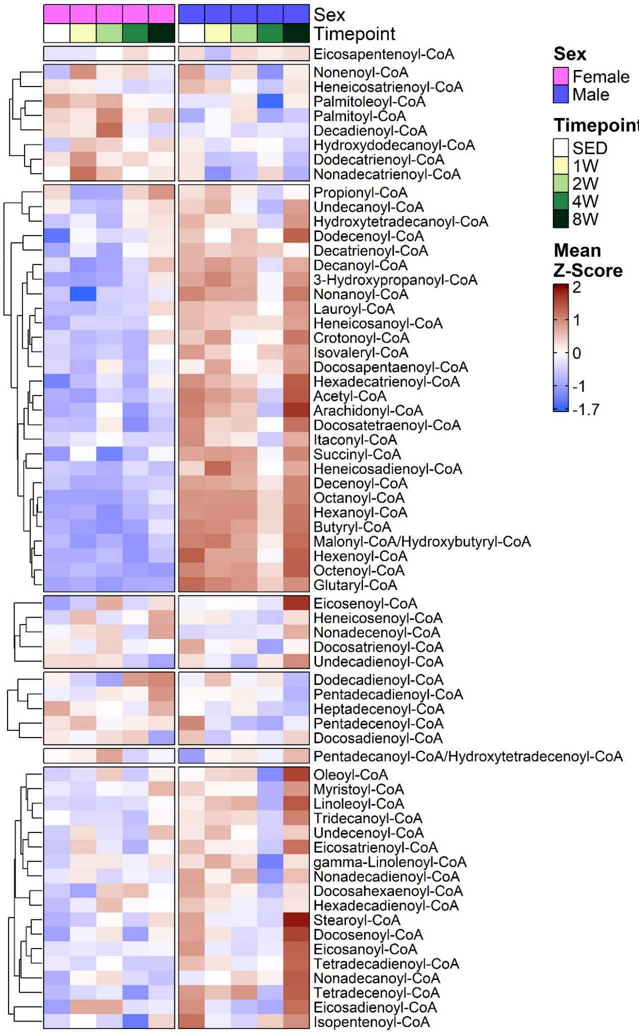


**B.** scWAT Amino Acids

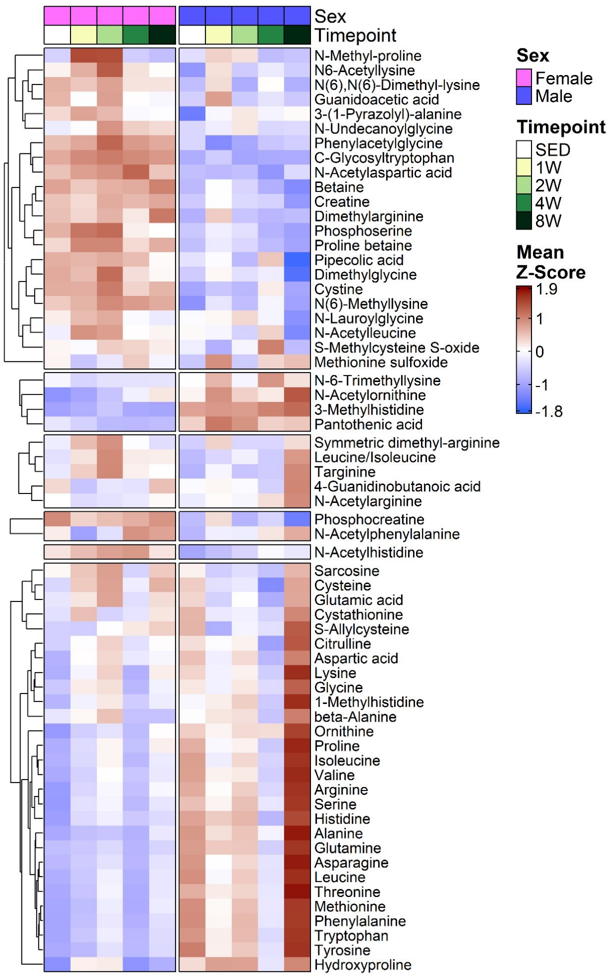


**C.** scWAT nucleotides

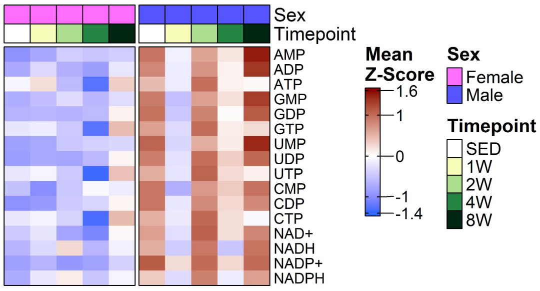

**Supplementary Figure S4. Metabolomic Visualizations.**

**A–C**) scWAT metabolome heatmaps displaying acyl-CoAs (**A**), amino acids (**B**), and nucleotides (**C**). Sample-level values were standardized before calculating the mean of each group to better observe patterns.

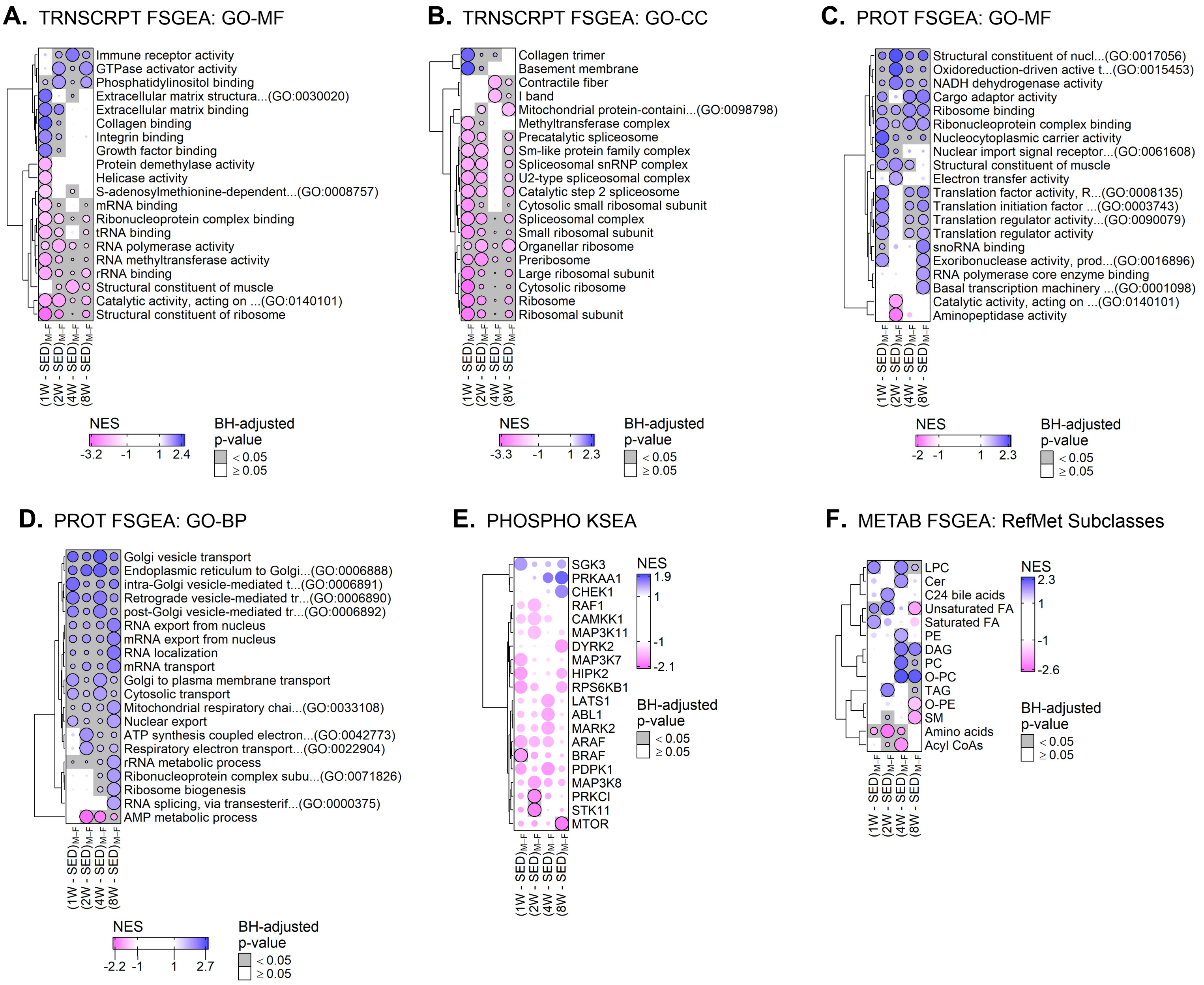
A. TRNSCRPT FSGEA: GO-MF
Immune receptor activity
GTPase activator activity
Phosphatidylinositol binding
Extracellular matrix structura...(GO:0030020)
Extracellular matrix binding
Collagen binding
Integrin binding
Growth factor binding
Protein demethylase activity
Helicase activity
S-adenosylmethionine-dependent...(GO:0008757)
mRNA binding
Ribonucleoprotein complex binding
tRNA binding
RNA polymerase activity
RNA methyltransferase activity
rRNA binding
Structural constituent of muscle
Catalytic activity, acting on ...(GO:0140101)
Structural constituent of ribosome
(1W - SED)M–F
(2W - SED)M–F
(4W - SED)M–F
(8W - SED)M–F
NES
-3.2
-1
1
2.4
BH-adjusted p-value
< 0.05
≥ 0.05
B. TRNSCRPT FSGEA: GO-CC
Collagen trimer
Basement membrane
Contractile fiber
I band
Mitochondrial protein-containi...(GO:0098798)
Methyltransferase complex
Precatalytic spliceosome
Sm-like protein family complex
Spliceosomal snRNP complex
U2-type spliceosomal complex
Catalytic step 2 spliceosome
Cytosolic small ribosomal subunit
Spliceosomal complex
Small ribosomal subunit
Organellar ribosome
Preribosome
Large ribosomal subunit
Cytosolic ribosome
Ribosome
Ribosomal subunit
(1W - SED)M–F
(2W - SED)M–F
(4W - SED)M–F
(8W - SED)M–F
NES
-3.3
-1
1
2.3
BH-adjusted p-value
< 0.05
≥ 0.05
C. PROT FSGEA: GO-MF
Structural constituent of nucl...(GO:0017056)
Oxidoreduction-driven active t...(GO:0015453)
NADH dehydrogenase activity
Cargo adaptor activity
Ribosome binding
Ribonucleoprotein complex binding
Nucleocytoplasmic carrier activity
Nuclear import signal receptor...(GO:0061608)
Structural constituent of muscle
Electron transfer activity
Translation factor activity, R...(GO:0008135)
Translation initiation factor ...(GO:0003743)
Translation regulator activity...(GO:0090079)
Translation regulator activity
snoRNA binding
Exoribonuclease activity, prod...(GO:0016896)
RNA polymerase core enzyme binding
Basal transcription machinery ...(GO:0001098)
Catalytic activity, acting on ...(GO:0140101)
Aminopeptidase activity
(1W - SED)M–F
(2W - SED)M–F
(4W - SED)M–F
(8W - SED)M–F
NES
-2
-1
1
2.3
BH-adjusted p-value
< 0.05
≥ 0.05
D. PROT FSGEA: GO-BP
Golgi vesicle transport
Endoplasmic reticulum to Golgi...(GO:0006888)
intra-Golgi vesicle-mediated t...(GO:0006891)
Retrograde vesicle-mediated tr...(GO:0006890)
post-Golgi vesicle-mediated tr...(GO:0006892)
RNA export from nucleus
mRNA export from nucleus
RNA localization
mRNA transport
Golgi to plasma membrane transport
Cytosolic transport
Mitochondrial respiratory chai...(GO:0033108)
Nuclear export
ATP synthesis coupled electron...(GO:0042773)
Respiratory electron transport...(GO:0022904)
rRNA metabolic process
Ribonucleoprotein complex subu...(GO:0071826)
Ribosome biogenesis
RNA splicing, via transesterif...(GO:0000375)
AMP metabolic process
(1W - SED)M–F
(2W - SED)M–F
(4W - SED)M–F
(8W - SED)M–F
NES
-2.2
-1
1
2.7
BH-adjusted p-value
< 0.05
≥ 0.05
E. PHOSPHO KSEA
SGK3
PRKAA1
CHEK1
RAF1
CAMKK1
MAP3K11
DYRK2
MAP3K7
HIPK2
RPS6KB1
LATS1
ABL1
MARK2
ARAF
BRAF
PDPK1
MAP3K8
PRKCI
STK11
MTOR
(1W - SED)M–F
(2W - SED)M–F
(4W - SED)M–F
(8W - SED)M–F
NES
1.9
1
-1
-2.1
BH-adjusted p-value
< 0.05
≥ 0.05
F. METAB FSGEA: RefMet Subclasses
LPC
Cer
C24 bile acids
Unsaturated FA
Saturated FA
PE
DAG
PC
O-PC
TAG
O-PE
SM
Amino acids
Acyl CoAs
(1W - SED)M–F
(2W - SED)M–F
(4W - SED)M–F
(8W - SED)M–F
NES
2.3
1
-1
-2.6
BH-adjusted p-value
< 0.05
≥ 0.05

**Supplementary Figure S5. Comparison of training response across sexes.**

**A–B)** Top molecular function (GO-MF) (**A**) or cellular component (GO-CC) (**B**) terms from the Gene Ontology database that are most significantly enriched in any of the 4 comparisons of the male and female training responses from the transcriptomics FGSEA results. **C–D)** Top GO-MF (**C**) or GO-BP (**D**) terms that are most significantly enriched in any of the 4 comparisons of the male and female training responses from the proteomics FGSEA results. **E)** Top kinases that are most significantly enriched in any of the 4 comparisons of the male and female training responses from the phosphoproteomics KSEA results. **F)** Top RefMet chemical subclasses that are most significantly enriched in any of the 4 comparisons of the male and female training responses from the metabolomics FGSEA results.

**A.** METAB Module Eigenfeatures

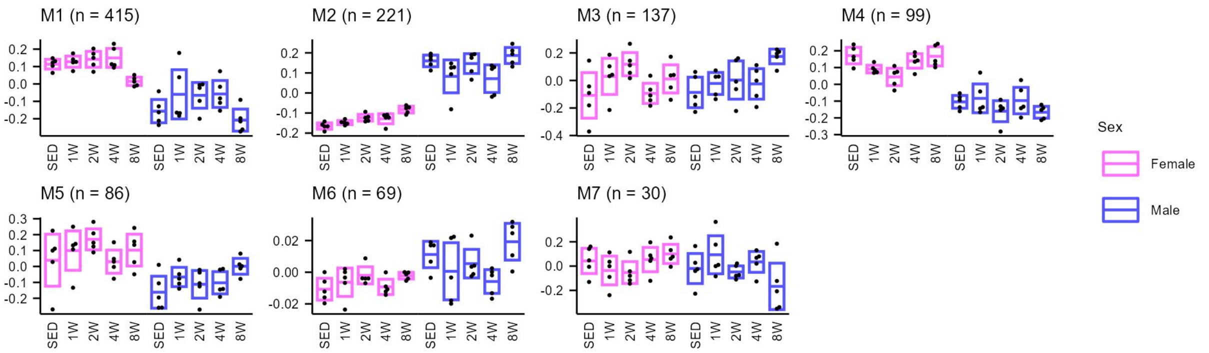


**B.** TRNSCRPT Module Eigenfeatures

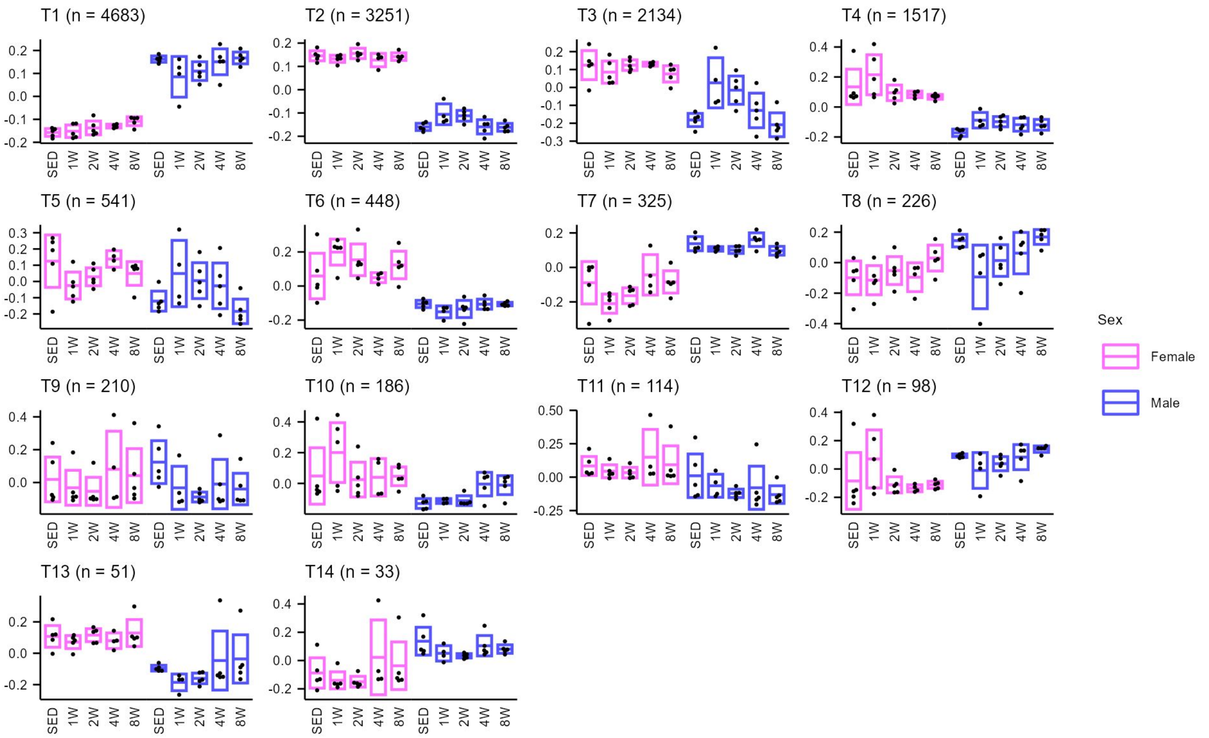


**C.** PROT Module Eigenfeatures

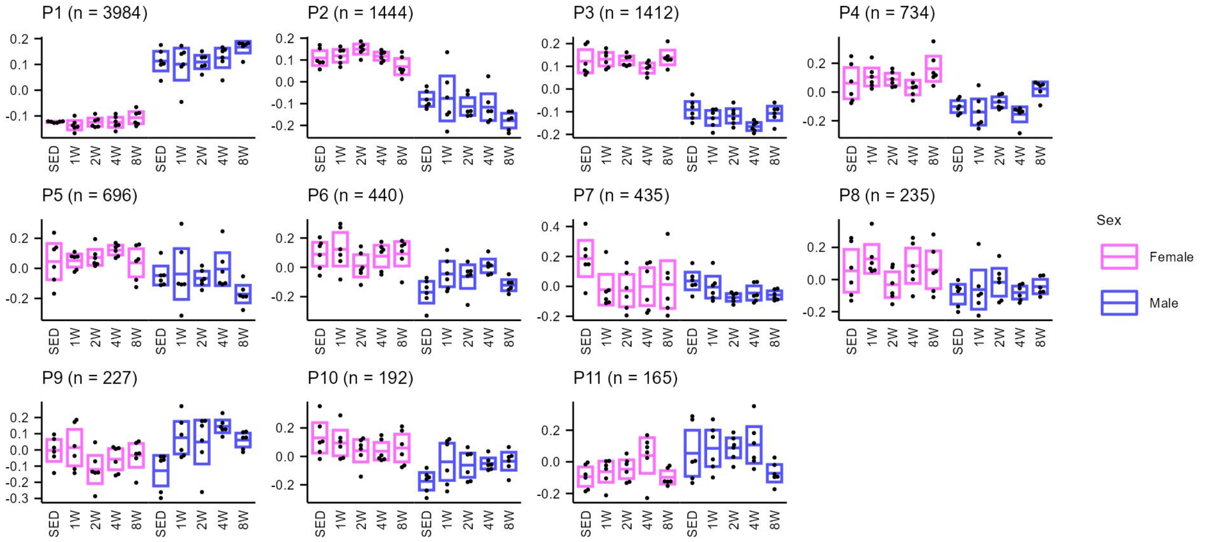

**Supplementary Figure S6. WGCNA module eigenfeatures.**

**A–C)** Plots of the module eigengenes (MEs) from the metabolomics (**A**), transcriptomics (**B**), and proteomics (**C**) WGCNA results. The size of each module is displayed along with the module labels. Boxes represent 95% confidence intervals for the mean of each group.

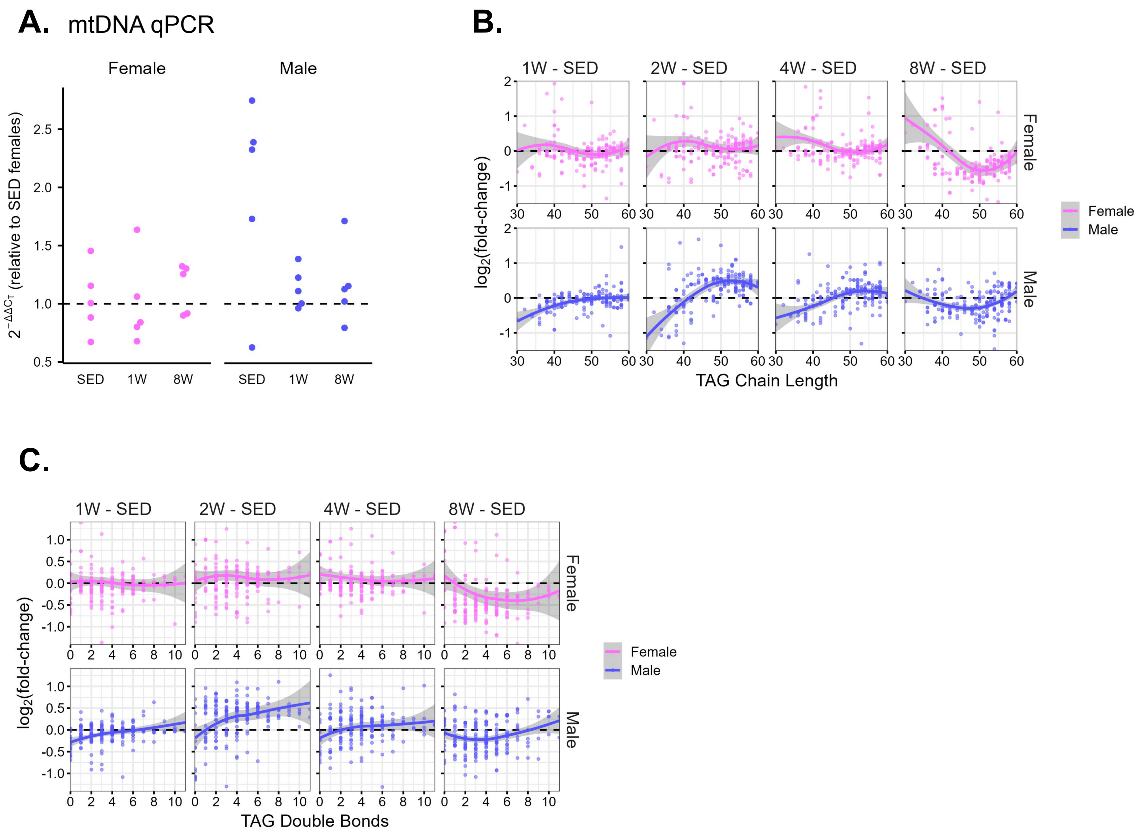
A. mtDNA qPCR
Female
Male
$2^{-\Delta\Delta C_T}$ (relative to SED females)
2.5
2.0
1.5
1.0
0.5
SED
1W
8W
SED
1W
8W
B.
1W - SED
2W - SED
4W - SED
8W - SED
Female
Male
$log_2$(fold-change)
TAG Chain Length
30
40
50
60
Female
Male
C.
1W - SED
2W - SED
4W - SED
8W - SED
Female
Male
$log_2$(fold-change)
TAG Double Bonds
0
2
4
6
8
10
1.0
0.5
0.0
-0.5
-1.0
Female
Male

**Supplementary Figure S7. Sexual dimorphism in lipid dynamics with ExT.**

**A)** mtDNA 2^-ΔΔ$C_T$ values as measured by real-time qPCR. **B–C)** Changes in chain length and double bond content of TAG species in male and female rats after 1, 2, 4, and 8 weeks of exercise training. Loess curves are included with 95% confidence bands. The y-axes have been restricted to more clearly show patterns, so not all points are visible.

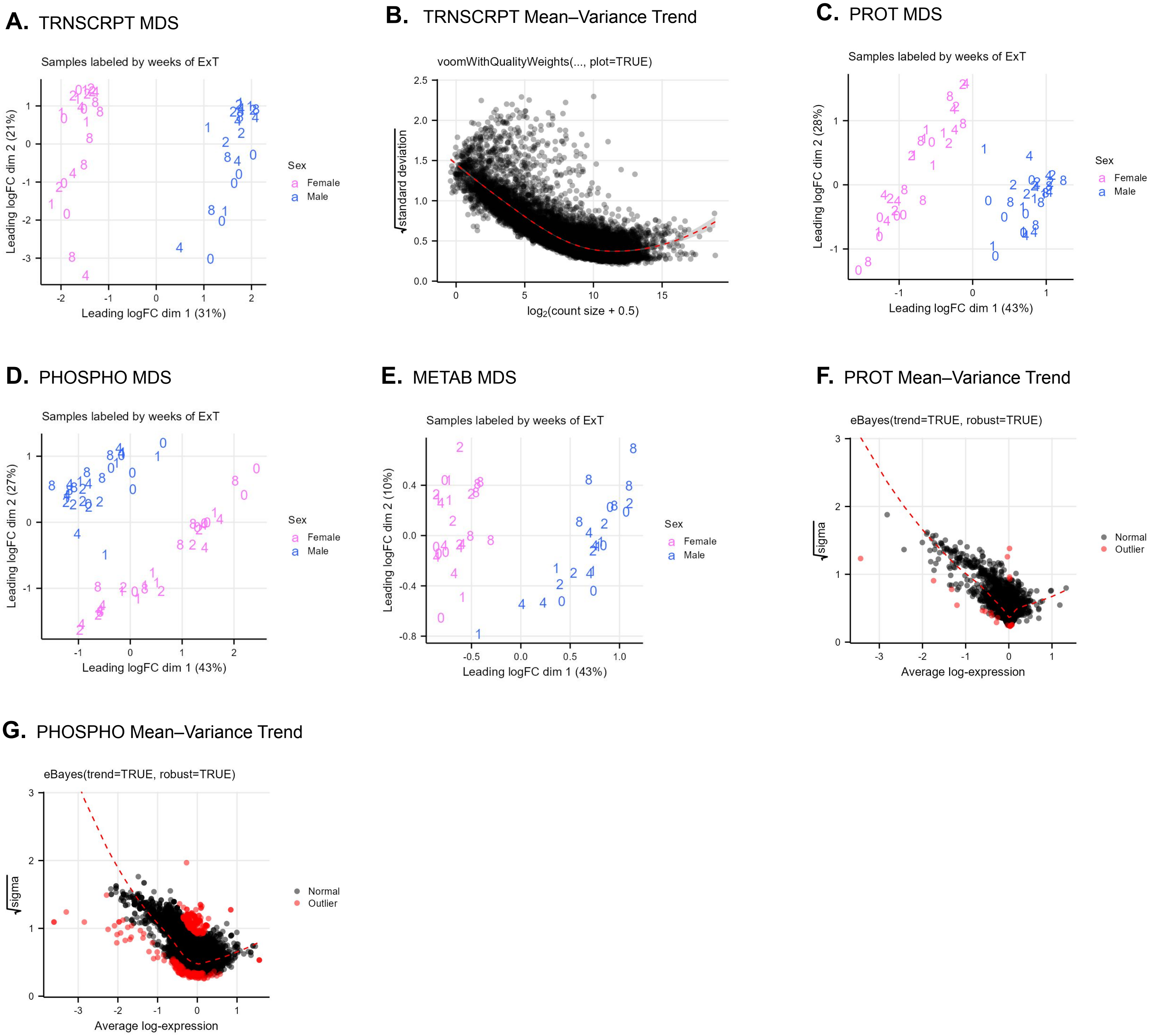

A. TRNSCRPT MDS
Samples labeled by weeks of ExT
Leading logFC dim 1 (31%)
Leading logFC dim 2 (21%)
Sex
Female
Male
B. TRNSCRPT Mean–Variance Trend
voomWithQualityWeights(..., plot=TRUE)
√standard deviation
log2(count size + 0.5)
C. PROT MDS
Samples labeled by weeks of ExT
Leading logFC dim 1 (43%)
Leading logFC dim 2 (28%)
Sex
Female
Male
D. PHOSPHO MDS
Samples labeled by weeks of ExT
Leading logFC dim 1 (43%)
Leading logFC dim 2 (27%)
Sex
Female
Male
E. METAB MDS
Samples labeled by weeks of ExT
Leading logFC dim 1 (43%)
Leading logFC dim 2 (10%)
Sex
Female
Male
F. PROT Mean–Variance Trend
eBayes(trend=TRUE, robust=TRUE)
√sigma
Average log-expression
Normal
Outlier
G. PHOSPHO Mean–Variance Trend
eBayes(trend=TRUE, robust=TRUE)
√sigma
Average log-expression
Normal
Outlier

**Supplementary Figure S8. Differential analysis model indicators.**

**A)** Multidimensional scaling (MDS) plots where distances between points approximate the typical $\log_2$ fold-changes between transcriptomics samples. **B)** Scatterplot of the average $\log_2$ counts per million vs. the square-root of the residual standard deviations from *limma::voom*. The loess curve (dashed red line) approximates the mean–variance relationship. **C–E)** MDS plots where distances between points approximate the typical $\log_2$ fold-changes between proteomics (**C**), phosphoproteomics (**D**), or metabolomics (**E**) samples. **F–G)** Average $\log_2$ relative abundance vs. the square-root of the residual standard deviations of proteins (**F**) or phosphosites (**G**). The dashed red line indicates the robust mean–variance trend from the *limma::eBayes* step. This is the *ggplot2* equivalent of the *limma::plotSA* output.
