## Supplementary Text for "Sexual dimorphism and the multi-omic response to exercise training in rat subcutaneous white adipose tissue"

### Supplementary Text: Data generation and processing

For complete detailed descriptions of methods used for sample preparation and multi-omics data generation and processing at chemical analysis sites for this study, please see the associated MoTrPAC publication[1].

##### *RNA-sequencing*

Total RNA was extracted from tissue lysates using a BiomekFx automation workstation. For blood samples, total RNA was extracted using the Agencourt RNAdvance blood specific kit (Beckman Coulter). RNA quantity and integrity was assessed with a Nanodrop (ThermoFisher Scientific, #ND-ONE-W), Qubit assay (ThermoFisher Scientific), and either Bioanalyzer or Fragment Analyzer. 500 ng of total RNA from each sample was used to generate libraries for RNA sequencing using the Universal Plus mRNA-Seq kit (NuGEN/Tecan #9133) and prepared with a Biomek i7 laboratory automation system (Beckman Coulter). Sequencing of pooled libraries was performed through 100bp paired-end sequencing using the Illumina NovaSeq 6000 platform (Illumina), targeting a sequencing depth of 35 millions read pairs per sample. Sequenced reads were demultiplexed with bcl2fastq2 (v2.20.0), adapters were trimmed with cutadapt (v1.18), pre-alignment QC metrics generated with FastQC (v0.11.8) and reads aligned using STAR (v2.7.0d). Quantification was performed using RSEM (v1.3.1).

##### *LC-MS/MS proteomics and phosphoproteomics*

Proteomics analyses were performed using clinical proteomics protocols described previously[2], with full details provided here[1]. scWAT samples were lysed and protein concentration was determined using BCA assay. Protein lysates were reduced with 5 mM dithiothreitol (DTT, Sigma-Aldrich) for 1 hour at 37°C with shaking at 1000 rpm on a thermomixer, alkylated with iodoacetamide (IAA, Sigma-Aldrich) in the dark for 45 minutes at 25°C with shaking at 1000 rpm, and diluted 1:4 with Tris-HCl, pH 8.0. Proteins were first digested with LysC endopeptidase (Wako Chemicals) at a 1:50 enzyme:substrate ratio (2 hours, 25°C, 850 rpm), followed by digestion with trypsin (Promega) at a 1:10 enzyme:substrate ratio (14 hours, 25°C, 850 rpm). Formic acid was added to a final concentration of 1% to quench digestion, after which peptides were desalted using Sep-Pac C18 columns (Waters) and BCA assay was used to determine final peptide concentrations.

Peptide aliquots (400 µg per sample) were resuspended to a final concentration of 5 µg/µL in 200 mM HEPES, pH 8.5 for isobaric labeling. Samples were randomized across the first 10 channels of tandem mass tag (TMT) 11-plexes (ThermoFisher Scientific), and the last channel (131C) of each multiplex was used for a common reference composed of a mix of peptides from all samples. TMT reagent was added to each sample at a 1:1 peptide:TMT ratio, and labeling proceeded for 1 hour at 25°C with shaking at 400 rpm. After labeling QC checks, reactions were quenched with hydroxylamine and samples within each multiplex were combined and desalted with Sep-Pac C18 columns (Waters). Each combined TMT multiplex was then fractionated using high pH reversed phase separation and concatenated into 24 fractions. 5% of each fraction was removed for global proteome analysis, and the remaining 95% was further concatenated to 12 fractions for phosphopeptide enrichment using immobilized metal affinity chromatography (IMAC).

For mass spectrometry analysis of the global proteome, online separation was performed using a nanoAcquity M-Class UHPLC system (Waters) and a 25 cm x 75 µm i.d. picofrit column packed in-house with C18 silica (1.7 µm UPLC BEH particles, Waters Acquity). Samples were analyzed with a Q Exactive HF mass spectrometer (ThermoFisher Scientific). For phosphoproteome samples, online separation was performed with a Dionex Ultimate 3000 UHPLC system (ThermoFisher Scientific) and a 30 cm x 75 µm i.d. picofrit column packed in-house with C18 silica (1.7 µm UPLC BEH particles, Waters Acquity). Samples were analyzed with a Q-Exactive HFX mass spectrometer (ThermoFisher Scientific). Full information regarding elution gradients and instrument settings for global proteomics and phosphoproteomics samples is described elsewhere[1].

Log2 TMT ratios to the universal reference were used as quantitative values for all proteomics features (full details of raw MS/MS data processing are described by the MoTrPAC Study Group[1]). Features not fully quantified in at least 2 TMT multiplexes and contaminant identifications were excluded from downstream analysis. Sample normalization was performed by median-centering and mean absolute deviation (MAD)-scaling within each sample, after which TMT multiplex batch effects were removed using the *limma::removeBatchEffect* function in R.

***Untargeted metabolomics***
Hydrophobic interaction liquid chromatography (HILIC) analyses of polar metabolites in the positive ionization mode were conducted at the Broad Institute of MIT and Harvard. 10 mg of cryopulverized tissue was homogenized in 300 µL of 10/67.4/22.4/0.018 v/v/v/v water/acetonitrile/methanol/formic acid containing stable isotope–labeled internal standards, and centrifuged at 9,000 x g for 10 min. Supernatants were injected onto a HILIC column (Waters) and analyzed using a Q-Exactive hybrid quadrupole Orbitrap mass spectrometer (ThermoFisher Scientific) operating in the positive mode. Raw data was processed for targeted peak integration using TraceFinder software (ThermoFisher Scientific) and with Progenesis QI software (Nonlinear Dynamics, Waters) for peak detection and integration of both metabolites with known identity and unknowns.

Reverse-phase and ion pairing profiling of polar metabolites was conducted at the University of Michigan. Non-pulverized tissue samples were weighed and homogenized in 1:1:1:1 methanol:acetonitrile:acetone:water (at a ratio of 1 mL per 50 mg tissue) using a sonicator. Samples were incubated on ice for 10 minutes followed by centrifugation at 15,000 x g for 10 minutes. Supernatant (300 µL) was dried in a nitrogen blower and reconstituted in water:methanol (8:2 v:v) for LC-MS analysis. Reverse phase analyses were performed on an Agilent 1290 Infinity II/ 6545 qTOF MS system with a Jetstream ESI source (Agilent Technologies) using a Waters Acquity HSS T3 column (Waters). Each sample was analyzed in both the positive and negative mode. Ion pairing analyses were performed on an identically-configured LC-MS system with an Agilent Zorbax Extend C18 1.8 µm RRHD column equipped with a matched guard column. Mass spectrometry analysis was conducted in the negative ion mode. Profinder v8.0 software (Agilent) was used for targeted compound detection and relative quantitation, while custom scripts were used for non-targeted feature detection. Agilent Mass Profiler Pro (v8.0) and Masshunter Qualitative Analysis were used for feature alignment and recursive feature detection. Features with >50% missing values across samples in a batch or >30% missing values in QC samples were removed, after which data reduction was performed using Binner[3] and normalized using the Systematic Error Removal Using Random Forest approach[4].

***Untargeted lipidomics***

For untargeted lipidomics analysis, 10 mg of tissue was homogenized in 400 µL isopropanol containing stable isotope-labeled internal standards from Avanti Polar Lipids (Alabaster) using freeze-thaw cycles in liquid nitrogen and sonication. Samples were centrifuged for 5 minutes at 21,000 x g and supernatants were used for LC-MS on a Vanquish chromatography system with an Accucore C30 column (2.1 x 150 mm, 2.6 µm particle size) coupled to a Q-Exactive HF mass spectrometer (ThermoFisher Scientific). Full details on elution gradients and instrument settings are provided elsewhere[1]. Raw LC-MS data was processed with Compound Discoverer v3.0 (ThermoFisher Scientific). Peak area was corrected for QC sample peak area across the batch and filtered with background and QC filters. Features absent in >50% of the QC pooled injections with a coefficient of variation <30% were removed from the dataset. Feature annotation was based on mass and relative abundance, retention time and MS2 patterns.

***Targeted metabolomics and lipidomics***

Branched-chain keto acids, acyl-CoAs and nucleotides were measured by targeted assays at Duke University. For analysis of branched-chain keto acids, 10 µL of plasma or 200 µL of tissue homogenate was extracted using ethyl acetate as previously described[5]. For Acyl-CoA extraction, 500 µL tissue homogenate was used as reported previously for liquid[6] and solid[7] phase extractions. Nucleotides were extracted as previously described[8,9]. Extractions were centrifuged at 14,000 x g for 5 minutes, and supernatants were analyzed by LC-MS/MS using a Xevo TQ-S triple quadrupole mass spectrometer (Waters). Endogenous levels were quantified by spiking tissue homogenates (acyl-coAs and nucleotides) or fetal bovine serum (keto acids) with authentic analytes (Sigma-Aldrich).

Amino acids and amino metabolites, TCA cycle metabolites, ceramides, and acylcarnitines were measured by targeted assays at the Mayo Clinic. For amino acid and amino metabolites, 5 mg of tissue homogenate was extracted and analyzed as previously described[10,11]. Ceramides and sphingolipids were extracted from 5 mg of tissue homogenate as previously described[12,13]; acylcarnitines were also extracted from 5 mg using previously described methods[14,15]. TCA metabolites were extracted from 5 mg of tissue and quantified using GC-MS[16], with minor modifications as reported elsewhere[1].

Targeted lipidomics analysis was performed at Emory University using 10 mg of cryopulverized tissue, homogenized in 100 µL PBS and diluted with 100 µL 20% methanol and spiked with 1% BHT solution according to previously used methods[17,18]. Homogenates were centrifuged at

14,000 x g for 10 minutes and supernatants were loaded onto C18 SPE columns and eluted with 400 µL methyl formate. External standards were purchased from Cayman Chemical. Samples were analyzed by LC-MS/MS using an ExionLC (SCIEX) chromatography system equipped with an Accucore™ C18 column (ThermoFisher) coupled to a SCIEX QTRAP 5500 mass spectrometer. Mobile phase A was water with 10 mM ammonium acetate, and mobile phase B was acetonitrile with 10 mM ammonium acetate. Details on the gradient program and subsequent mass spectrometry analysis are provided elsewhere[1]; raw data was processed using Sciex OS (AB SCIEX, v1.6.1).

***Metabolomics data processing and normalization***

All metabolomics datasets were partitioned into named compounds (for analytes that were confidently identified) and unnamed compounds (for those without a standard chemical name). Only named metabolites were included in this analysis. Data was $\log_2$-transformed, and analytes with >20% missing values were removed. For targeted datasets with >12 analytes, and for all untargeted datasets, missing values were imputed using K-nearest neighbors (k=10). Median sample–sample correlation was used to identify outlier samples, which were manually reviewed by the metabolomics sites. Untargeted datasets were normalized using sample median-centering. For details on the removal of redundant measures of analytes detected on multiple platforms, please refer to the Supplementary Note associated with this manuscript.
